# Nitrogen nutrition impacts grapevine esca leaf symptom incidence, physiology and metabolism

**DOI:** 10.1101/2024.08.31.610625

**Authors:** Ninon Dell’Acqua, Gregory A. Gambetta, Gwenaëlle Comont, Nathalie Ferrer, Adam Rochepeau, Pierre Pétriacq, Chloé E. L. Delmas

## Abstract

Nitrogen plays a crucial role in plant growth and defence mechanisms, yet its role in plant-pathogen interactions is complex and remains largely unexplored, especially in perennial crops. This study aimed to investigate the effects of controlled nitrogen nutrition levels on disease incidence, fungal communities, and plant physiology and metabolism. Esca is a widespread grapevine vascular disease affecting physiology, xylem integrity and metabolism. Naturally infected *Vitis vinifera* L. cv. Sauvignon blanc were subjected to three ammonium nitrate treatments across three seasons, resulting in reduced esca incidence under nitrogen deficiency compared with medium nutrition levels, while excess nitrogen had no significant impact. Nitrogen treatments significantly impacted vine physiology and leaf metabolites but did not affect fungal wood communities. Nitrogen deficiency significantly reduced stem diameter, photosynthesis, and leaf area, likely decreasing whole-plant transpiration, while excess nitrogen increased these factors suggesting a key role of plant transpiration in esca incidence. Additionally, nitrogen deficiency led to significantly higher production of phenylpropanoids, particularly flavonoids, in leaf metabolomes compared to the medium level. These findings highlight the pivotal role of nitrogen in the development of esca through alterations in vine morphology, physiology and metabolism. Fertilization practices may be crucial in the management of plant diseases.

## INTRODUCTION

Nitrogen is the most important nutrient that plants take up from the soil and, after water, is the main limiting factor for plant growth (Vitousek and Howarth, 1991). Nitrogen plays a crucial role in both the development and defence of plants, having a wide range of effects on physiological and biochemical processes, and plant-pathogen interactions (Fagard *et al*., 2014). Plants can often experience nitrogen-stress conditions due to either nitrogen deficiency or nitrogen excess from past and present fertilizer applications. Although viticulture is considered as a crop with low fertilization demands compared to annual crops, nitrogen still plays an important role in producing quality wine (Jackson and Lombard, 1993). Managing nitrogen in vineyards is complex as it is an equilibrium between controlling vine vigor, maximizing grape quality, mitigating environmental impact and managing fertilization costs (Verdenal *et al*., 2021). Nitrogen deficiency induces a slowdown in vine growth resulting in a reduction in leaf area, a lower number of leaves on secondary shoots, and faded green leaf color (Metay *et al*., 2015; Vrignon-Brenas *et al*., 2019). Nitrogen deficit also changes the biomass allocation leading to biomass reduction in annual organs in favour of perennial ones, such as trunks (Metay *et al*., 2015). Nitrogen excess, on the contrary, leads to a high vigour with a dense canopy and large dark green leaves (Verdenal *et al*., 2021).

Plant growth rate is one of the main traits influenced by nitrogen status as it mostly relies on the production of primary metabolites, enzymes, and structural proteins, which are synthesized from carbon assimilation during photosynthesis. The functioning of the photosynthetic apparatus depends on nitrogen availability, as up to 75% of the total nitrogen content of leaves is used in photosynthesis (Seepaul *et al*., 2016) underpinned by highly abundant enzymes like Rubisco and thylakoid proteins (Ferrario-Méry *et al*., 1997). Physiological responses involved in CO_2_ assimilation such as stomatal conductance, net photosynthesis, and transpiration could therefore be impacted by nitrogen levels (Zhang *et al*., 2021). A correlation has been shown between nitrogen fertilization and CO_2_ assimilation in grapevines (Keller *et al*., 2001; Schreiner *et al*., 2013), except in Squeri *et al*. (2021). This relationship may depend on plant age, localisation nitrogen form or genotypes. One assumption made in the literature is that, during moderate to strong nitrogen deficit, plant growth would be reduced to a greater extent than CO_2_ assimilation leading to a carbohydrate over-accumulation (Herms and Mattson, 1992). This nitrogen-carbon imbalance would lead to a shift from primary metabolism to the production of secondary metabolites, including the accumulation of defence compounds which could impact immune response (Liao *et al*., 2022). Integrated studies that combine plant health, growth, physiological, and metabolomic measurements could facilitate the evaluation of nitrogen’s potential impact on defence metabolism (Massad *et al*., 2012), and consequently, on the potential influence of nitrogen in plant-pathogen interaction.

Plant resistance to disease is also influenced by nitrogen levels (Huber and Watson, 1974), but depending on multiple factors such as concentration (Danial and Parlevliet, 1995), type of nitrogen fertiliser (Mur *et al*., 2017), time of application (Abuley *et al*., 2019), plant species (Mundy, 2008; Soulie *et al*., 2020) and type of pathogens (Fagard *et al*., 2014). The effect of tissue nitrogen concentration on plant susceptibility appears to depend on the pathogen species. In the case of tomato, there was no effect on susceptibility to *Fusarium oxysporum* (a hemibiotrophic pathogen), but susceptibility to *Pseudomonas syringae* (also hemibiotrophic) and *Oidium lycopersicum* (a biotrophic pathogen) increased significantly with nitrogen concentration (Hoffland *et al*., 2000). However, a decrease in susceptibility to *Botrytis cinerea* (a necrotrophic pathogen) with increasing nitrogen concentration has been shown (Hoffland *et al*., 1999). Finally, plants are likely to be more susceptible to biotrophic pathogens and more resistant to necrotrophic pathogens when fertilized with nitrogen (Dordas, 2008; Snoeijers *et al*., 2000).

In grapevines, very little research has been done on nitrogen-plant-pathogen interactions. Most of the research has been done on *Botrytis cinerea*, *Erysiphe necator*, and *Plasmopara viticola* (Bavaresco and Eibach, 1987; Keller *et al*., 2003; Mundy, 2008; Marcianò *et al*., 2023), and demonstrated that pathogen severity increased with nitrogen supply. Studies have suggested that nitrogen may influence vascular disease by altering plant physiology, but no specific studies have been carried out so far (Destrac-Irvine *et al*., 2007; Calzarano *et al*., 2009). Among vascular diseases in vineyards, esca remains one of the most problematic because there is still no cure available (Claverie *et al*., 2020). Esca is a complex disease associated with foliar symptoms, most commonly found in grapevines between 10 and 30 years old, depending on the cultivar (Etienne *et al*., 2024; Gastou *et al*., 2024). Foliar symptoms are observed randomly from one year to the next (Dewasme *et al*., 2022) and are not reproducible by inoculating the fungal pathogens responsible for wood degradation and necrosis in the trunk (Mugnai *et al*., 1999). Thus, esca expression can only be studied in the field or using potted uprooted mature plants that are naturally infected and express leaf symptoms under controlled conditions with the same probability as in the field (Bortolami et al. 2021; Dell’Acqua et al. 2023). Recent studies have suggested that low grapevine vigor is correlated with low esca incidence in the field (Gastou et al. 2024; Claverie et al. 2025). However, the impact of different nitrogen fertilization levels on esca expression is still unknown. Nitrogen, which can influence water use efficiency, growth, and hydraulic traits, is likely to have a direct impact on esca pathogenesis. Recent works have highlighted the role of water transport within the plant and the influence of varietal water use efficiency on the development of esca leaf symptoms (Bortolami *et al*., 2021*b*; Gastou *et al*., 2024). A decrease in whole plant stomatal conductance and the reduction in *CO_2_* assimilation, stomatal conductance and *WUE_i_* at symptomatic leaf level has been demonstrated in two grapevine varieties (Bortolami *et al*., 2021*b*; Dell’Acqua *et al*., 2023). An alteration in the radial stem growth during esca pathogenesis has also been recently reported (Bortolami *et al*., 2021*b*; Dell’Acqua *et al*., 2023). Esca is also known to impact grapevine leaf and fruit metabolomes (Lima *et al*., 2017; Moret *et al*., 2020; Ouadi *et al*., 2021; Weiller *et al*., 2024). Furthermore, esca is associated with different fungal pathogens and many of them are hemibiotrophic, i.e. organisms that live within living tissue for part of their lives and then continue to live in dead tissue (Bruez *et al*., 2014, 2016, 2020; Elena *et al*., 2018). On one hand, pathogen community composition could influence pathogen response to nitrogen fertilization. On the other hand, nitrogen fertilization could influence wood pathogen abundance, diversity and activity and thus esca pathogenesis.

This study aimed to investigate the effects of grapevine nitrogen nutrition on vascular pathogenesis while monitoring plant growth, physiology and metabolism, and trunk fungal communities. We hypothesized that under nitrogen deficit, the reduction in plant transpiration and the production of secondary metabolites would drive a decrease in leaf symptom incidence. Conversely, an excess of nitrogen could increase the incidence of esca leaf symptoms by enhancing plant growth and resource availability for pathogen development. We studied esca pathogenesis under different nitrogen conditions, using naturally infected mature *Vitis vinifera* cv. Sauvignon blanc vines, transplanted from the field into pots and subjected to three different levels of nitrogen (in the form of ammonium nitrate) over three vegetative seasons. Finally, to further explore the effect of nitrogen nutrition on grapevine susceptibility to wood pathogens, we inoculated rooted cuttings produced from the three nitrogen levels and compared the length of necrosis caused by two grapevine trunk pathogen species.

## MATERIALS AND METHODS

### Plant material

The experiment was realized over three vegetative seasons (2021, 2022 and 2023) using 99 *Vitis vinifera* cv. Sauvignon blanc vines aged of ∼30-y-old, 81 grafted onto 101-14 Millardet et de Grasset (101-14 MGt) uprooted from a vineyard planted in 1992 at INRAE Nouvelle-Aquitaine Bordeaux (44°47024.800N, 0°34035.100 W) in 2017, 2018 and 2019 and 18 grafted onto Fercal and uprooted from a vineyard planted in 1991 at Couhins near Bordeaux (44°45ʹ17.9′′N, 0°33ʹ32.6′′W) in 2021. The presence of esca leaf symptoms had been monitored in these vineyards at the plant level since 2012, following Lecomte *et al*. (2012). Plants were uprooted during late winter and transferred into 20L pots as described in Bortolami *et al*. (2019). Transplantation is the only method allowing the study of esca foliar symptom development under controlled conditions as it is only observed on mature plants in the field. Esca expression in these mature potted plants occurred naturally at the same rate as in the field under well-watered conditions and with similar severity (Bortolami et al. 2021; Dell’Acqua et al. 2024). Plants were pruned to retain four one-bud spurs on each of the two arms (and eight one bud spurs on one arm if there was only one remaining). The plants were installed in a polytunnel and distributed in six rows: two for each nitrogen level with an equivalent proportion of uprooting years and rootstocks. In 2021 and 2022, there were 32 plants in the lowest nitrogen level ([NH_4_NO_3_]_low_), 33 plants in the middle level ([NH_4_NO_3_]_medium_), and 34 plants in the highest nitrogen level ([NH_4_NO_3_]_high_). During the 2022 season, two plants died : one in the low nitrogen level and the other in the medium level. Climate (i.e. humidity (in %) and temperature (in °C)) was monitored and from these parameters maximal vapor pressure deficit (*VPD*) was calculated for the three years of experiments (Supplementary Fig. S1). Images of the plants before transplantation and during the experiment are presented in Supplementary Fig. S2.

### Fertilisation management

Three different nutritive solutions were prepared using varying concentrations of NH_4_NO_3_ (0.25 mM, 2 mM, and 8 mM for the [NH_4_NO_3_]_low_, [NH_4_NO_3_]_medium_, and [NH_4_NO_3_]_high_ treatment, respectively) and, for all solutions, KH_2_PO_4_ (0.4 mM), CaCl_2_ (0.75 mM), MgSO_4_. 7H_2_O (0.5 mM), K_2_SO_4_ (0.3 mM), and oligoelements. Plants were continuously drip-irrigated to ensure the absence of water stress (mean predawn water potential < -0.3MPa, data not shown). To maintain distinct plant nitrogen levels throughout the four-month period across the three nitrogen treatments, we conducted weekly monitoring of the plants using a N-tester. The N-tester optically measures the chlorophyll content of the leaf, which is related to the nitrogen status of the plant (Evans et al. 1989). A N-tester index value is obtained for a series of 30 measurements (i.e. non-destructive leaf pinches). For each nitrogen level and date, three values were obtained, using leaves of similar age and exposition. If the plants from [NH_4_NO_3_]_low_ exhibited a N-tester index that was too similar to [NH_4_NO_3_]_medium,_ they were irrigated with the same nutritive solution as described above, but with [NH_4_NO_3_] = 0 until the N-tester index differed. The same method was applied for [NH_4_NO_3_]_high_ in comparison to [NH_4_NO_3_]_medium_. The mean N-tester index values over the season of [NH_4_NO_3_]_low,_ [NH_4_NO_3_]_medium_ and [NH_4_NO_3_]_high_ (Supplementary Fig. S3) were aligned with very low, medium and very high N-tester index identified in the literature (Van Leeuwen and Friant, 2011). In addition, in the second and third years (ie. 2022 and 2023), we monitored plants of the same cultivar (Sauvignon blanc) in an INRAE vineyard closeby the polytunnel with no nitrogen nutrition problems (VITADAPT experimental block; (Destrac and van Leeuwen, 2016) to manage the fertilization of the [NH_4_NO_3_]_medium_ level according to the field N-tester index (Supplementary Fig. S3) and the [NH_4_NO_3_]_low_ and [NH_4_NO_3_]_high_ as the first year. Images of the experiment are presented in Supplementary Fig. S2. To compare nitrogen level scores between the three treatments, we performed a linear mixed model per year with the day of year set as a random effect and the nitrogen level treatment as a fixed effect. A post-hoc Tukey test adjustment for multiple comparisons was realized. We confirmed statistically that this method resulted in significantly higher and lower nitrogen levels compared to medium level for each three years (*P* value < 0.001 for each treatment comparison; Table S1). As we managed the [NH_4_NO_3_]_medium_ level according to the vineyard, we did not expect to have significantly the same values for the [NH_4_NO_3_]_medium_ level and the vineyard. However, in 2022 there was no significant difference in N-tester index between the [NH_4_NO_3_]_medium_ level and the vineyard (*P* value =0.056; Table S1).

### Esca leaf symptom notation

Leaf symptom expression was monitored weekly over a four month-period at the leaf and canopy levels during all three years of the experiment. Esca leaf symptoms (Supplementary Fig. S2) were observed between July 19 and September 7, 2021, June 2 and September 7, 2022 and between June 16 and August 22, 2023. During a season, entire plants could be noted as asymptomatic (hereafter called control, when none of the leaves were symptomatic) or symptomatic (when at least 25% of the canopy presented tiger-stripe leaf symptoms). We also scored symptoms at the leaf scale according to 3 classes: “C”, asymptomatic leaf from control plants, “S” symptomatic leaf from symptomatic plants and “AS”, asymptomatic leaf from symptomatic plants (both before and after symptom appearance), as done by Bortolami *et al*. (2021*a*).

To compare leaf symptom expression between nitrogen levels, we performed a linear mixed model with the year set as a random effect and the nitrogen level treatment as a fixed effect. A post-hoc Tukey test adjustment for multiple comparisons was realized.

### Vigor, stomatal density and gas exchange measurements

#### Plant growth measurements

Plant growth was assessed in control plants in each nitrogen level early September 2022 using (i) stem diameter at the third internode measured at 1 cm above the bottom node on the dorso-ventral axis, and (ii) the average length of three secondary shoots (taken from the sixth, seventh, and eighth nodes). These traits were measured in 20 control plants in [NH_4_NO_3_]_low_,19 plants in [NH_4_NO_3_]_medium_, and 18 plants in [NH_4_NO_3_]_high_.

At the end of the season on November 30, 2023, the senescence stage was scored in control plants (n=86) and symptomatic plants (n=13) using an extended version of the Biologische Bundesanstalt, Bundessortenamt und Chemische Industrie (BBCH) phenological scale adapted for grapevine (Lorenz *et al*., 1995). This system is a useful tool that provides a code for similar stages of development.

#### Leaf area

To estimate plant vigor, we calculated an average leaf area for each nitrogen level in control plants. On September 19, 2022, the area of approximately 100 healthy mature leaves from the middle of the canopy was measured per nitrogen level using a LI-3000, LI-COR.

#### Stomatal density analysis

We used five leaves per nitrogen level to estimate stomatal density in control plants. As stomata are mainly located on the under leaf surface in grapevines, nail polish was applied under the leaf blade to the same location and left to dry for 30 minutes. The imprint was removed with transparent scotch tape. Three images were then taken for different localisations on each leaf impression, avoiding veins as much as possible, under a binocular magnifying glass with a Nikon SMZ1270 camera, and the NIS-ElementsD software associated with the magnifier. Each picture was analysed with imageJ by counting each stomata for the three localisations per leaf. The stomatal density was then assessed by dividing the number of stomata by the surface picture area in mm^2^.

#### Leaf gas exchange

Ambient stomatal conductance (*g_s_*) was measured with a Li-600 (LICOR) on control leaves between April 26 and September 9, 2022. Measurements were taken on control plants between 10h and 12h on 11 dates, on a total of 282 control leaves in the [NH_4_NO_3_]_low_ level, 250 leaves in [NH_4_NO_3_]_medium_ , and 289 leaves in [NH_4_NO_3_]_high_. To compare the ambient stomatal conductance of symptomatic and control plants before symptom expression, we retrospectively selected measurements of “AS” leaves (from plants that became symptomatic later in the season) and “C” leaves (from control plants) taken in April and May 2022, on six dates. We thus used measurements from 13 leaves in [NH_4_NO_3_]_low_ level, 41 leaves in [NH_4_NO_3_]_medium,_ and 25 leaves in [NH_4_NO_3_]_high_.

During symptom expression between July 6 and September 9, 2022 (5 dates), ambient stomatal conductance of symptomatic leaves were measured in 8 “S” leaves from 2 symptomatic plants in [NH_4_NO_3_]_low_, 25 “S” leaves from 7 symptomatic plants in [NH_4_NO_3_]_medium_, and 19 “S” leaves from 4 symptomatic plants,,in [NH_4_NO_3_]_high_.

Gas exchange was measured in the 2022 season between 9h and 12h on mature well-exposed leaves from the middle of the stem of control asymptomatic plants using the TARGAS-1 portable photosynthesis system (PP Systems). Cuvette Photosynthetic Active Radiation was set to optimize photosynthesis (1500 μmol m^−2^ s^−1^). The light-saturated photosynthetic rate (*Amax* in µmol m^−2^ s^−1^) and the maximum stomatal conductance (*g_smax_* in mmol m^−2^ s^−1^) was recorded by alternating rows of plants (i.e. nitrogen levels). Measurements were performed at four dates between July and September 2022 with a total of 35 leaves in the [NH_4_NO_3_]_low_ level, 56 leaves in [NH_4_NO_3_]_medium_ and 38 leaves in [NH_4_NO_3_]_high_.

The water use efficiency (*WUE_i_*) was calculated as the ratio between light-saturated photosynthetic rate (*Amax* in µmol m^−2^ s^−1^) and maximum stomatal conductance (*g_smax_* in mmol _m-2 s-1)._

#### Statistical analyses

To compare vigor (dorso-ventral stem diameter, lateral shoot length, and leaf area), stomatal density, and gas exchange measurements (*g_s_*, *g*_smax_, *A*_max_ and *WUE*_i_) of control asymptomatic plants, between the three nitrogen levels, we performed ANOVAs, associated with a post-hoc tukey test after normality was checked with a Levene test.

### Trunk fungal communities

#### Sampling method

The trunk sampling was carried out in apparently healthy wood on September 21, 2022 using a drill at the level of the graft union. For each level of nitrogen fertilization, 10 control vines were sampled (for a total of n= 30 plants) and all the esca-symptomatic vines (n= 2, 7, and 4 plants in [NH_4_NO_3_]_low_ , [NH_4_NO_3_]_medium_ and [NH_4_NO_3_]_high_, respectively) resulting in 43 plants sampled. For each vine, a 2cm^2^ window in the bark was removed at the grafting point using a sterilized razor blade, then cleaned with ethanol. If the wood was apparently healthy and free of necrotic lesions, wood pieces were collected in a petri dish by drilling to a maximum depth of 1.5 cm using a drill (Worx Power, Positec Germany) and a 0.7 mm drill bit cleaned between each vine. This depth likely corresponds to functional wood in mature grapevines (McElrone *et al*., 2021) and is also the type of wood where a xylem discolored stripe developed during esca pathogenesis (Lecomte *et al*., 2024). We therefore hypothesize that this sampling site is a critical location where pathogenic activity can occur. When necrotic wood was reached with the drill, a new location was chosen as close as possible to the previous one. The wood pieces were placed in 2.5 mL tubes with a sterilized tweezer and directly immersed in liquid nitrogen. The samples were then placed at -80°C until grinding by a tissue-lyser (TissueLyser II, Qiagen, Germany).

#### DNA extraction

For each sample (n=43), 60 mg was weighed and placed into a 2 ml tube in liquid nitrogen. Aliquots of 60 mg are stored at -80°C until extraction. We used the DNeasy Plant Mini Kit (Qiagen, Germany) to extract the total genomic DNA according to the manufacturer’s protocol. All samples were diluted to obtain a maximum of 15 ng/µL of DNA. A negative extraction control was included in the process.

#### Fungal ITS1 amplification and Illumina MiSeq sequencing

Two successive PCRs are used to amplify the targeted region and attach sample-specific adapters and indexes on either side. The internal transcribed spacer 1 (ITS1) region of the fungal ITS ribosomal DNA gene (Schoch *et al*., 2012) was amplified with the ITS1F-ITS2 primers. Universal sequences were added to the specific sequences to enable hybridization of adapters and sequencing indexes during 2nd PCR (i.e. mb-**ITS1F:** 5‘-TCGTCGGCAGCGTCAGATGTGTATAAGAGACAG**CTTGGTCATTTAGAGGAAGT AA**-3’ and mb-**ITS2:** 5’-GTCTCGTGGGCTCGGAGATGTGTATAAGAGACAG**GCTGCGTTCTTCATCGATG C**-3’, with the specific sequences in bold).

The first PCR mixture (final volume = 25μl) consisted of 12.5μl of 2X mix PCR (Platinum™ Hot Start PCR Master Mixes, Invitrogen, Thermo Fisher Scientific Inc., USA), 5.5µL eau DNase/RNase free, 2.5μl each of the forward and reverse primers and 2μL of DNA template. PCR was performed using a thermocycler (Mastercycler EP Gradient, Eppendorf, Germany) under the following conditions: initial denaturation at 95°C for 2 minutes; followed by 35 cycles of 95°C for 45 s, 55°C for 60 s, and 72°C for 90 s; and a final extension phase at 72°C for 10 min. Two negative PCR controls were included in the analysis. These PCR controls were subjected to the same amplification procedure as all of the other samples used to rule out potential contaminants detected by metabarcoding. The quality of the amplification products was checked by electrophoresis in 2% agarose in TBE 0.5X gels with 5µL of PCR product and 2µL of Zvision (2X) and migration for 30min at 100V.

The second PCR and the amplicon sequencing (Illumina V3 kit: 2 × 300 bp paired-end reads) were performed by the Genome Transcriptome Facility of Bordeaux according to standard protocols.

#### Bioinformatics and diversity analyses

The bioinformatic analyses were carried out using the FROGS tools hosted by the Galaxy platform (Escudié *et al*., 2018). After obtaining the fastq sequences, the FROGS preprocessing tool was used to associate R1 and R2 and obtain unique sequences. Then the clustering, chimera removal, OTU (Operational Taxonomic Unit) filter, ITS sequence selection and affiliation tools (blast on the database ITS UNITE Fungi 8.3), of the FROGS software were used to obtain the abundance table. The data curation was made using the MetabaR package in R (Zinger *et al*., 2021) by taking into account the extraction and PCR controls (Table S2). When necessary, OTUs were blasted on a database specific to grapevine trunk disease pathogens, trunkdiseaseID.org (Table S3). The MetabaR package was then used to transform the clean data obtained to an object readable by the R phyloseq package v1.34.0 (McMurdie and Holmes, 2013).

In order to evaluate the impact of nitrogen level on the wood fungal communities, control plants (n= 30) were used to calculate alpha and beta diversity indices. The observed, Shannon, and Simpson alpha diversity indices of the trunk fungal communities were compared between nitrogen levels using Kruskal Wallis tests. To visualize the dispersions among samples, we performed ordination plots using a non-metric multidimensional scaling (NMDS) ordination matrix based on Bray-curtis distance. Beta dispersion was calculated from transformed centered log-ratio (CLR) data to test the overall differences in composition between nitrogen level groups in control plants using a PERMANOVA. We compared alpha and beta diversity between control plants (n=10 for each nitrogen level) and symptomatic plants, only in the medium and high nitrogen levels (n=4 and 7 symptomatic plants, respectively), as only two plants were symptomatic in the low nitrogen level and thus excluded from the analysis (reducing overall sample size to 41 plants).

To find associations between microbial features, nitrogen level, and symptoms, we performed general linear models using MaAsLin2 (Mallick *et al*., 2021). Fungal communities analyses were performed using the phyloseq package in R (McMurdie and Holmes, 2013) and MicrobiomeAnalyst 2.0 (Lu *et al*., 2023).

### Untargeted metabolomic analysis

#### Sampling method

The wood samples from control plants (n=30) used for metabarcoding were also used for untargeted metabolomic analysis (10 mg of lyophilized wood samples).

Green leaves were sampled from randomly selected control plants (i.e. 17 plants in [NH_4_NO_3_]_low_, 13 plants in [NH_4_NO_3_]_medium_ and 17 plants in [NH_4_NO_3_]_high_) during the 2022 season in the different nitrogen nutrition levels and at different dates, always in the morning (between 8h and 11h). They were selected on the primary shoots, at the same position and exposure and directly placed in liquid nitrogen and then stored at -80°C until further analysis.

#### Metabolomics of semi-polar extracts

We extracted semi-polar compounds, including primary and secondary metabolites, using automated high-throughput ethanol extraction procedures at the MetaboHUB-Bordeaux Metabolome (https://metabolome.u-bordeaux.fr/) from 10 mg of freeze-dried grapevine leaves and wood material, following previously established protocols (Luna *et al*., 2020). Untargeted metabolic profiling by UHPLC-LTQ-Orbitrap mass spectrometry (LCMS) was performed as already reported (Dussarrat *et al*., 2022; Martins *et al*., 2022) using an Ultimate 3000 ultra-high-pressure liquid chromatography (UHPLC) system coupled to an LTQ-Orbitrap Elite mass spectrometer interfaced with an electrospray ionisation source (ESI, ThermoScientific, Bremen, Germany) operating in negative ion modes as described previously (Luna *et al*., 2020).

All 66 leaves (i.e. 20 leaves from [NH_4_NO_3_]_low_, 22 leaves from [NH_4_NO_3_]_medium_ and 24 leaves from [NH_4_NO_3_]_high_) samples and 30 wood samples (i.e. 10 samples per nitrogen level treatments) runs for the LCMS analytical sequence were randomised, extraction blanks (prepared without plant material and used to rule out potential contaminants detected by untargeted metabolomics), and Quality Control (QC) samples that were prepared by mixing 20 µL from each sample. QC samples were injected every 10 runs and used for i) the correction of signal drift during the analytical batch, and ii) the calculation of coefficients of variation for each metabolomic feature so only the most robust ones are retained for chemometrics (Broadhurst *et al*., 2018). Briefly, MS1 full scan acquisitions at high-resolution (240k) were performed on QC samples and the various standards for exact mass measurements, and all samples were subjected to MS2 Data Dependent Analysis (DDA, 30k resolution) to generate fragmentation information for further annotation (Martins *et al*., 2022).

#### Processing of metabolomic data

Raw LCMS data were processed using MS-DIAL v 4.9 (Tsugawa *et al*., 2015). We also implemented retention time correction in MS-DIAL, and the metabolomic signals were normalised according to the LOWESS method based on the QC samples. We performed data-curation according to blank check and setting thresholds as signal-to-noise ratio (SN) > 10, coefficient of variation in quality controls < 30%. The 3184 final features for predictive metabolomics annotations were performed using the MetaboHUB AgroMix in-house chemical database and the FragHUB database (Dablanc *et al*., 2024). Thus, putative annotation of differentially expressed metabolites resulted from MS-DIAL screening of the MS1 detected exact HR m/z and MS2 fragmentation patterns (Tsugawa *et al*., 2015). Additionally, the InChiKeys of annotated features were employed within ClassyFire to automatically generate a structural ontology for chemical entities (Djoumbou Feunang *et al*., 2016). Putative metabolites that were unlikely to be found in leaves and woods (e.g. Human drugs) were considered as misassigned and therefore reassigned as “Unknown”.

### Statistical analysis

Normalization to sample median, square root data transformation and Pareto scaling were applied to metabolomic data. We performed principal component analysis (PCA) for leaf and trunk samples to compare the metabolome of control plants between dates and between nitrogen levels. Volcano plots were performed to identify biomarkers between treatments. Thresholds were set at *P* value < 0.05 (Welch’s t-test) and a twofold change threshold at FC > 1.5 using a Benjamini–Hochberg correction for false-discovery rate (FDR). Statistical analysis was performed using Metaboanalyst and R.

#### Effect of nitrogen fertilization on grapevine susceptibility to wood pathogens

During vine dormancy (winter of 2022), we sampled woody cuttings from the Sauvignon blanc plants grown under the three different levels of fertilization. Cuttings were planted in March 2023 in 0.5 l individual pots filled with Klass-man RHP 15 commercial potting mix [fair peat of sphaine (70%), cold black peat (15%), pearlite and Danish clay (15%)], and were incubated for six weeks in a greenhouse (25°C, 16 h light/8 h dark) in Villenave d’Ornon (INRAE, France). After 3 months, the cuttings were inoculated with *Phaeomoniella chlamydospora* (strain CBS 239.74) and *Neofusicoccum parvum* (strain VIE35). These two isolates are part of the culture collections of INRAE (UMR SAVE, Villenave d’Ornon, France) and are stored at 4°C. Prior to inoculation, we replicated the fungal strains at 22°C on malt agar (w:w, 3 :4) in Petri dishes. To inoculate the cuttings, 5mm diameter wounds were made on each vine using a drill. The wounds were filled with a mycelium plug cut from the margin of each fresh mycelial culture and sealed with paraffin film (Parafilm M, Bemis, USA). A total of 15 cuttings were inoculated for each fungus and nitrogen level, with 5 cuttings per nitrogen level inoculated with non-colonised malt-agar plugs as negative controls (mock). Three months after inoculation, the cuttings were cut longitudinally and the size of the internal necrosis measured and compared between nitrogen levels using a non-parametric Kruskal-Wallis test.

## RESULTS

### Nitrogen levels impacted esca incidence

During the three years of monitoring, we observed esca leaf symptoms in 24.75% of the plants (n=99) at least once a year, which is typical for Sauvignon Blanc of this age in the plots of origin. Among those 25 plants, two expressed esca symptoms each of the three years, 10 expressed symptoms in two among the three years, and 13 plants expressed in only one of the three years. After three years of experiment, we found that plants under nitrogen deficiency exhibited significantly fewer esca leaf symptoms than those with regular nitrogen nutrition (i.e. 6.4% ± 0.1 (± se) of symptomatic vines in [NH_4_NO_3_]_low_ level vs. 19.5% ± 5.6 in [NH_4_NO_3_]_medium_ level; Fig. 1). The incidence of esca leaf symptoms under [NH_4_NO_3_]_high_ was 13.7% ± 2.0 on average, and did not significantly differ from the two other levels (Fig. 1).

**Figure 1.**
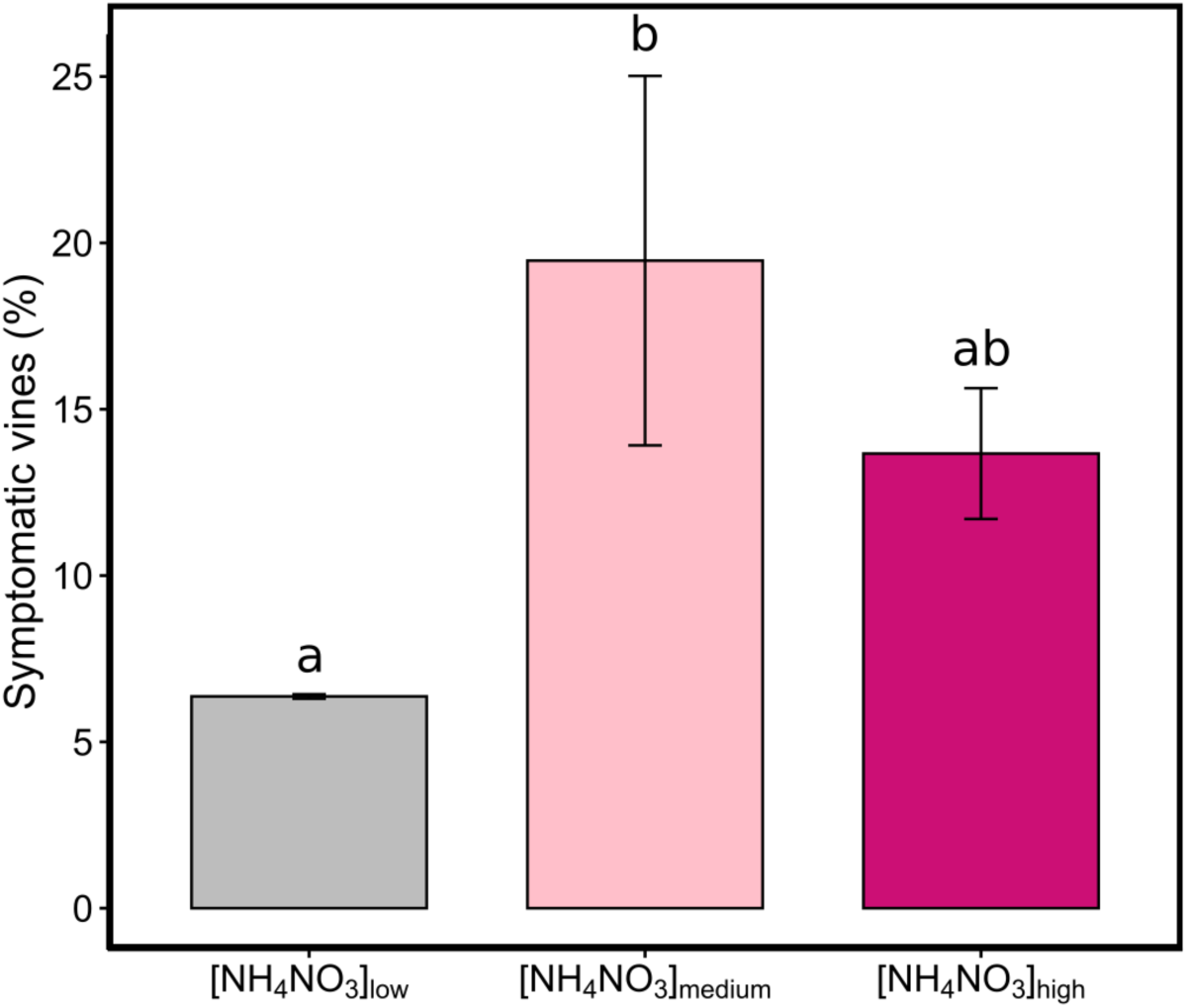
Mean percentage of esca-symptomatic vines *(Vitis vinifera* cv. Sauvignon blanc) according to the three levels of nitrogen nutrition during the three experimental seasons. Low nitrogen level ([NH_4_NO_3_]_10_w) is represented in grey, medium nitrogen level ([NH_4_NO_3_]medium) in pink and high nitrogen level ([NH_4_NO_3_]high) in dark pink. Error bars are standard errors. A linear mixed-effects model with the year set as a random effect and the nitrogen level treatment as a fixed effect was used. Letters represent significant differences (P value < 0.05, post-hoc tukey test).

### Vine growth, leaf characteristics and senescence differed under different nitrogen levels in control plants

The vegetative growth of control plants was significantly impacted by nitrogen nutrition. Stem diameter, secondary shoot length, and leaf area significantly increased with the nitrogen level (*P* value < 0.05, ANOVA, Tukey post hoc, Fig. 2A,B,C). On the contrary, there was no significant difference in stomatal density between the different nitrogen levels (Fig. 2D).

**Figure 2.**
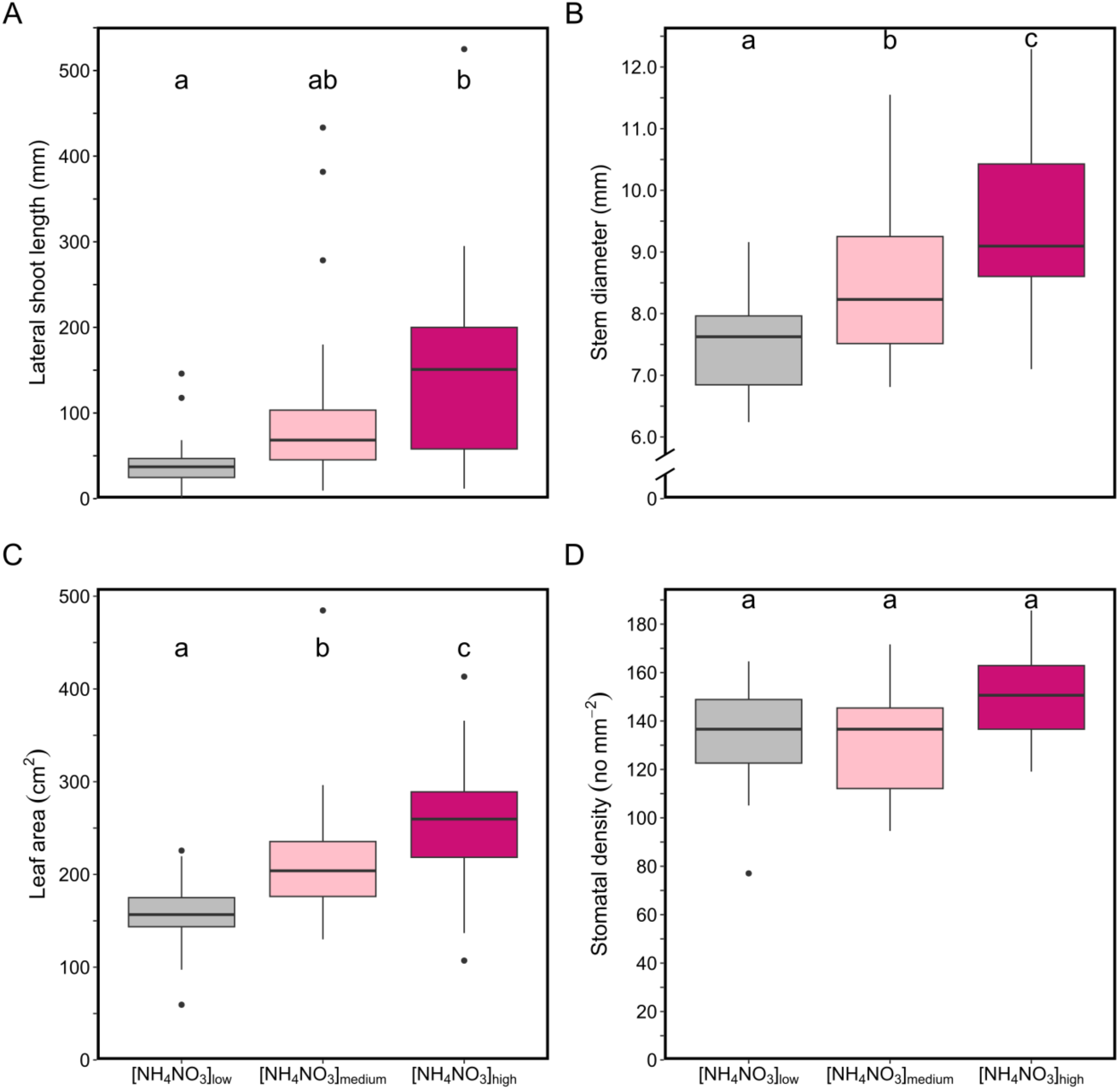
Growth and structural traits of *Vitis vinifera* cv. Sauvignon blanc plants subjected to three different nitrogen nutrition levels in 2022. (A) Lateral shoot length (in mm) and (B) dorso-ventral stem diameter (in mm) were measured on 20 plants in [NH_4_NO_3_]_10_w, 19 plants in [NH_4_NO_3_Jmedium, and 18 plants in [NH_4_NO_3_]high, (C) leaf area of -100 leaves from each nitrogen level (in cm^2^), (D) leaf abaxial stomata! density (5 leaves per nitrogen level, in no mm-^2^). Boxplots display the median and interquartile range, with whiskers extending to the minimum and maximum values, excluding outliers, which are shown as individual black points.The horizontal line in the box represents the median value. The letters indicate significant differences in ANOVAs (P value < 0.05, post-hoc tukey test).

On November 30, 2023, 100% of the leaves of control vines had fallen in level [NH_4_NO_3_]_low_ (i.e.100% of control vines were stage 97 on the BBCH scale), 40% of the vines had lost at least 50% of their leaves in [NH_4_NO_3_]_medium_ level (i.e. 40% of control vines were at least stage 95 on the BBCH scale) and none of the leaves had fallen in [NH_4_NO_3_]_high_ level (i.e. 100% of control vines didn’t have reached stage 93 on the BBCH scale) (Supplementary Fig. S4). The same shift in senescence under low and high nitrogen level was observed in 2022 but plant phenology was not monitored. We observed that when the symptomatic plants managed to produce new green shoots, they exhibited delayed senescence compared to the control plants, regardless of the nitrogen level treatment.

### Gas exchange in control plants differed under different nitrogen levels

The ambient stomatal conductance measurements in control leaves evolved throughout the 2022 vegetative season and significantly differed between nitrogen levels for several dates during the season (*P* value < 0.05, ANOVAs, Supplementary Fig. S5). There was no difference in ambient stomatal conductance between green leaves from control and future symptomatic plants (data not shown). During symptom expression, ambient stomatal conductance of symptomatic leaves was 23.8 ± 4.22 (se) mmol m^−2^ s^−1^ on average compared to 97 ± 3.8 (se) mmol m^−2^ s^−1^ on average in control plants and did not vary significantly between nitrogen levels (*P* value=0.3, ANOVA).

In control leaves, maximum CO_2_ assimilation (*A_max_*) was significantly lower under [NH_4_NO_3_]_high_ or [NH_4_NO_3_]_low_ levels compared to [NH_4_NO_3_]_medium_ level (*P* value < 0.05, Fig. 3A). The maximum stomatal conductance (*gs_max_*) under [NH_4_NO_3_]_low_ level did not significantly differ from [NH_4_NO_3_]_medium_ level. On the contrary, the *gs_max_* was significantly reduced under [NH_4_NO_3_]_high_ level (*P* value < 0.05, Fig. 3B). Consequently, the water use efficiency at leaf scale (*WUEi*) increased significantly with nitrogen levels (*P* value < 0.05, ANOVA, Tukey post hoc, Fig. 3C).

**Figure 3.**
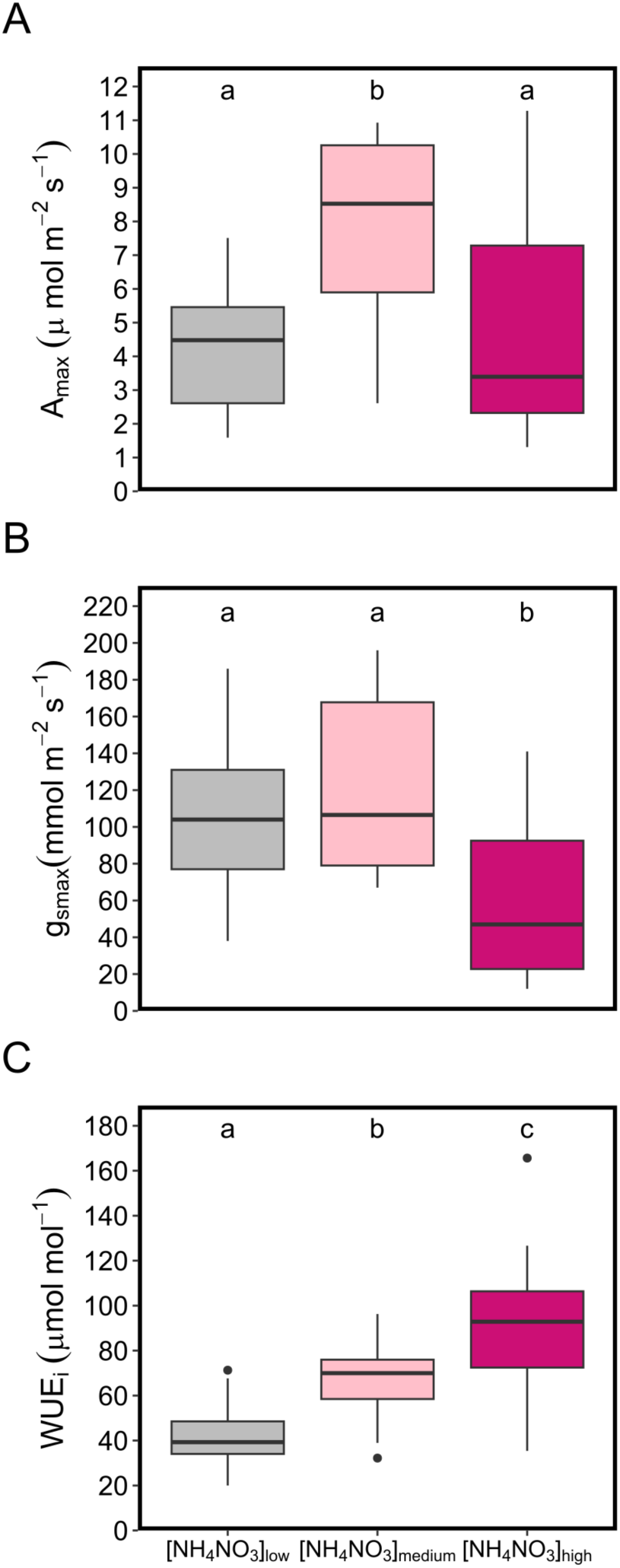
Leaf physiological traits of Vitis vinifera cv. Sauvignon blanc plants subjected to three different nitrogen nutrition levels during the 2022 season. (A) light-saturated photosynthetic rate (Amax in µmol m-^2^ s-1), (B) maximum stomata! conductance (gsmax, in mmol m-^2^ s-1), (C) water use efficiency (WUE;, in µmol m-^2^ s-^1^). The measurements were made on 35 control leaves in [NH_4_NO_3_]_10_w, 56 leaves in [NH_4_NO_3_lmedium, and 38 leaves in [NH_4_NO_3_]high· The letters indicate significant differences in ANOVAs (P value < 0.05, post-hoc tukey test). Boxplots display the median and interquartile range, with whiskers extending to the minimum and maximum values, excluding outliers, which are shown as individual black points.

### Trunk fungal communities in varying nitrogen levels and esca-symptomatic status

After paired-end alignments and cleaning process 2,019,933 fungal ITS sequences were generated from 41 samples, with an average of 49,266 fungal ITS sequences per sample (Table S2). These reads have been assigned to 337 Operational Taxonomic Unit (OTU) as presented in Table S3. Raw read number per sample and OTU are presented in Table S4 (including the two samples from the two symptomatic plants under low nitrogen level excluded from further analysis).

In healthy wood samples, the 20 most abundant OTUs were identified as belonging to the following genus: *Acremonium, Ambrosiella, Atrocalyx, Cladosporium, Devriesia, Filobasidium, Fomitiporia, Neofusicoccum, Penicillium, Phaeoacremonium*, and *Phaeomoniella.* In relative abundance, the proportions of these genus were found to vary depending on the sample (Fig. 4).

**Figure 4.**
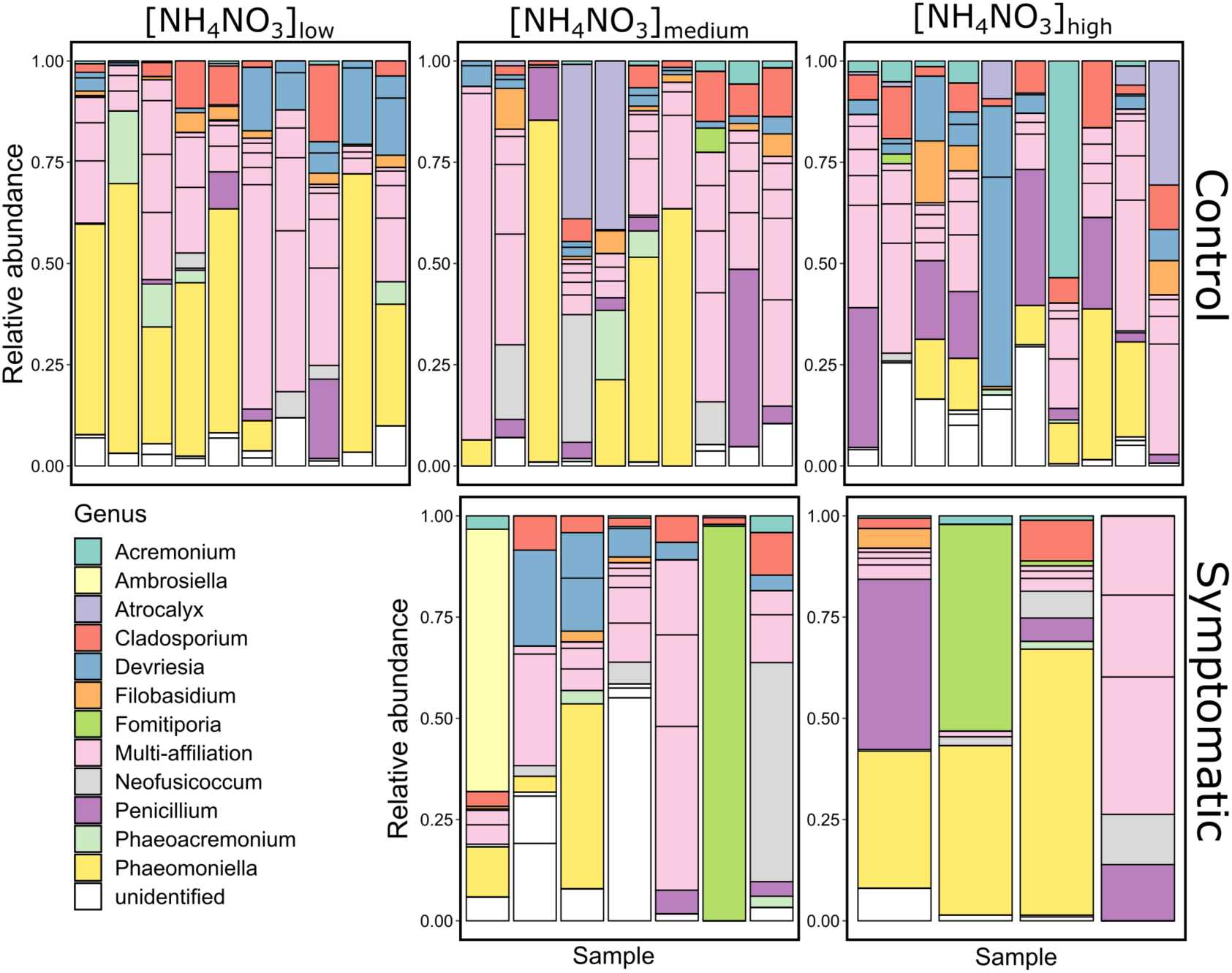
Relative abundance of the top 20 fungal Operational Taxonomic Unit (OTU) classified by genus in control and symptomatic *Vitis vinifera* cv. Sauvignon blanc trunk samples (healthy wood) from the three nitrogen levels. Each bar represents one sample. Note the absence of samples from symptomatic plants in the [NH_4_NO_3_]_10_w level, as only two plants were symptomatic (not included).

The main fungal species associated with grapevine trunk diseases in the literature (i.e. *Anthostoma decipiens, Botryosphaeria dothidea, Botryosphaeria stevensii, Cadophora luteo-olivacea, Diaporthe ampelina, Diplodia seriata, Eutypa lata, Eutypa sp, Fomitiporia mediterranea, Lasiodiplodia theobromae, Neofusicoccum parvum, Phaeoacremonium minimum, Phaeomoniella chlamydospora, Phellinus punctatus, Pleurostoma richardsiae, and Stereum hirsutum*) were found partly in almost all healthy wood samples from both control and symptomatic plants (except for two control samples collected from the high nitrogen level) (Supplementary Fig. S6).

### Diversity indexes of fungal microbiota were not significantly affected by nitrogen level

Regarding alpha diversity (the diversity within samples), the fungal communities were not significantly different between nitrogen treatments in control plants from the 2022 season: observed diversity, Shannon (i.e. specific diversity) and Simpson (measuring the probability that two randomly selected individuals belong to the same species) indexes were not significantly different (*P* value > 0.05, Kruskal wallis tests, Fig. 5). Regarding beta diversity (i.e. the measurement of community dissimilarity), NMDS indicated no significant differences either between control plants (*P* value > 0.05, PERMANOVA, Supplementary Fig. S7). Using multiple linear regressions, we found that none of the species could be used as nitrogen level biomarkers.

**Figure 5.**
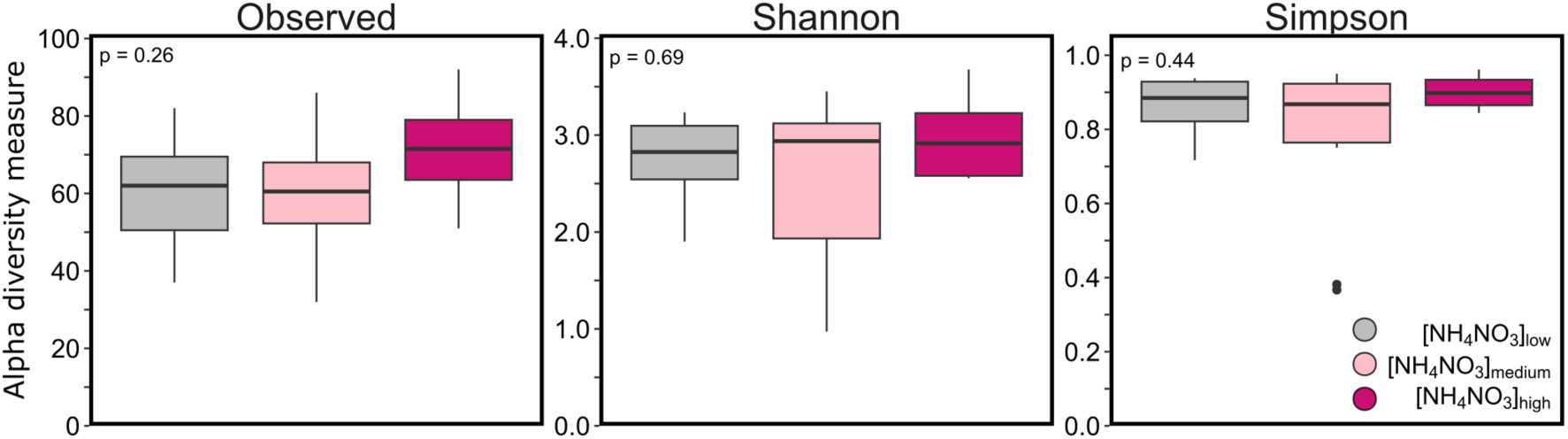
Observed, Shannon, and Simpson alpha diversity indices of the trunk fungal communities in healthy wood of control plants under three nitrogen levels. No significant differences were found between treatments (P value > 0.05, Kruskal Wallis test). Boxplots display the median and interquartile range, with whiskers extending to the minimum and maximum values, excluding outliers, which are shown as individual black points.

### Diversity indexes of fungal microbiota were partly affected by symptom expression

When comparing the alpha diversity indexes of wood fungal microbiota from control and the few symptomatic plants collected, we found no significant differences in [NH_4_NO_3_]_medium_ level, while the Shannon and Simpson indexes were significantly lower in symptomatic plants in [NH_4_NO_3_]_high_ level (*P* value < 0.05, Wilcoxon test, Fig. 6A). In addition, we found a significant difference in beta diversity when comparing symptom-nitrogen level groups (*P* value < 0.05, PERMANOVA, Fig. 6B).

**Figure 6.**
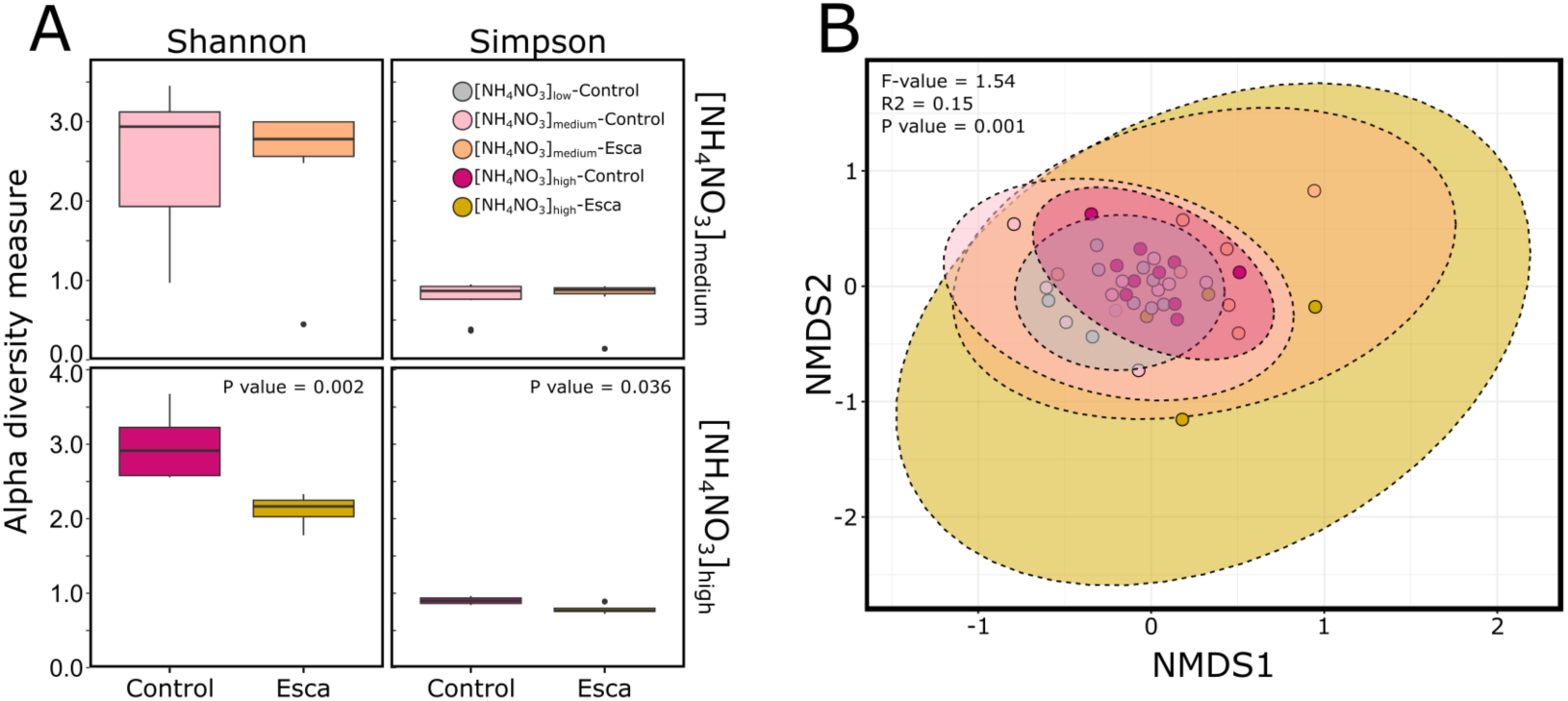
Analysis of fungal communities from healthy wood between control and symptomatic *Vitis vinifera* cv. Sauvignon blanc and nitrogen level treatments. (A) Shannon and Simpson alpha diversity indices, (B) Beta diversity. Five groups were compared in the analysis: [NH_4_NO_3_]_10_w - Control (n=10, in grey), [NH_4_NO_3_lmedium - Control (n=10, in pink), [NH_4_NO_3_lmedium - Esca (n=7, in orange), [NH_4_NO_3_]high - Control (n=10, in dark pink) and [NH_4_NO_3_]high - Esca (n=4, in yellow). Significant differences have been found between treatments in alpha diversity measurements (only P value < 0.05 are indicated in the figure, Kruskal Wallis test). Boxplots (A) display the median and interquartile range, with whiskers extending to the minimum and maximum values, excluding outliers, which are shown as individual black points. Beta dispersion (B) was calculated from transformed centred log-ratio (CLR) data to test the difference in dissimilarity between nitrogen level groups using a PERMANOVA (F value = 1.53, R^2^ = 0.15, P value = 0.001).

Using general linear models to search for associations between microbial features and symptomatology, pooling medium and high nitrogen levels, no fungi associated with esca in the literature were identified as significant biomarkers.

### Leaf and trunk metabolomes varied between nitrogen levels, especially phenylpropanoid metabolites

The leaf metabolism of control plants evolved over the course of the 2022 season (Supplementary Fig. S8). Consequently, we decided to separate the control samples in two time periods: early summer period, from June to the beginning of July 2022 and summer period, from the end of July to the beginning of August 2022. For both periods, PCA showed clear differences between the three nitrogen levels for control leaves (Fig. 7A,B). In order to study the metabolome associated with early symptom development, we focused on the early summer period.

**Figure 7.**
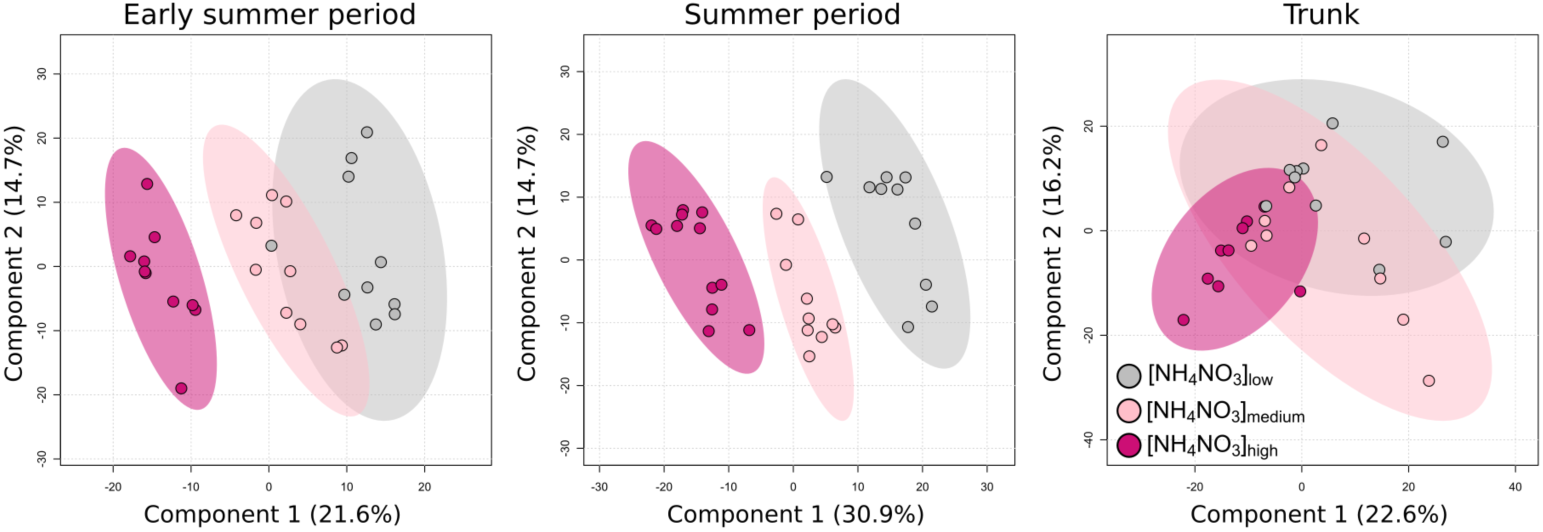
Principal component analysis (PCA) of metabolic profiles in leaves and trunks of *Vitis vinifera* cv. Sauvignon blanc. PCAs compare the three nitrogen levels in control leaves collected during (A) the early summer period (June 7, 2022 June 24, 2022 and July 4, 2022) and (B) the summer period (July 21, 2022, August 10, 2022), and (C) trunk samples collected on September 12, 2022. Colored circles arbitrarily enclose treatment types: low nitrogen level [NH_4_NO_3_]_10_w (in grey), medium nitrogen level [NH_4_NO_3_lmedium (in pink), and high nitrogen level [NH_4_NO_3_]h,gh (in dark pink).

Comparing the [NH_4_NO_3_]_low_ level to the [NH_4_NO_3_]_medium_ level of the early summer period, we found 47 metabolomic features significantly enriched in the low nitrogen level and 14 features significantly depleted (Fig. 8A). Putative metabolites annotated as primary metabolites, such as amino acids (e.g. glutamic acid), carbohydrates, and lipids (e.g. saccharolipids) were depleted in [NH_4_NO_3_]_low_ level (Fig. 8B) while secondary metabolites, such as phenylpropanoids and benzenoids were enriched. The phenylpropanoids enriched in [NH_4_NO_3_]_low_ were mainly flavonoids (Fig. 8C).

**Figure 8.**
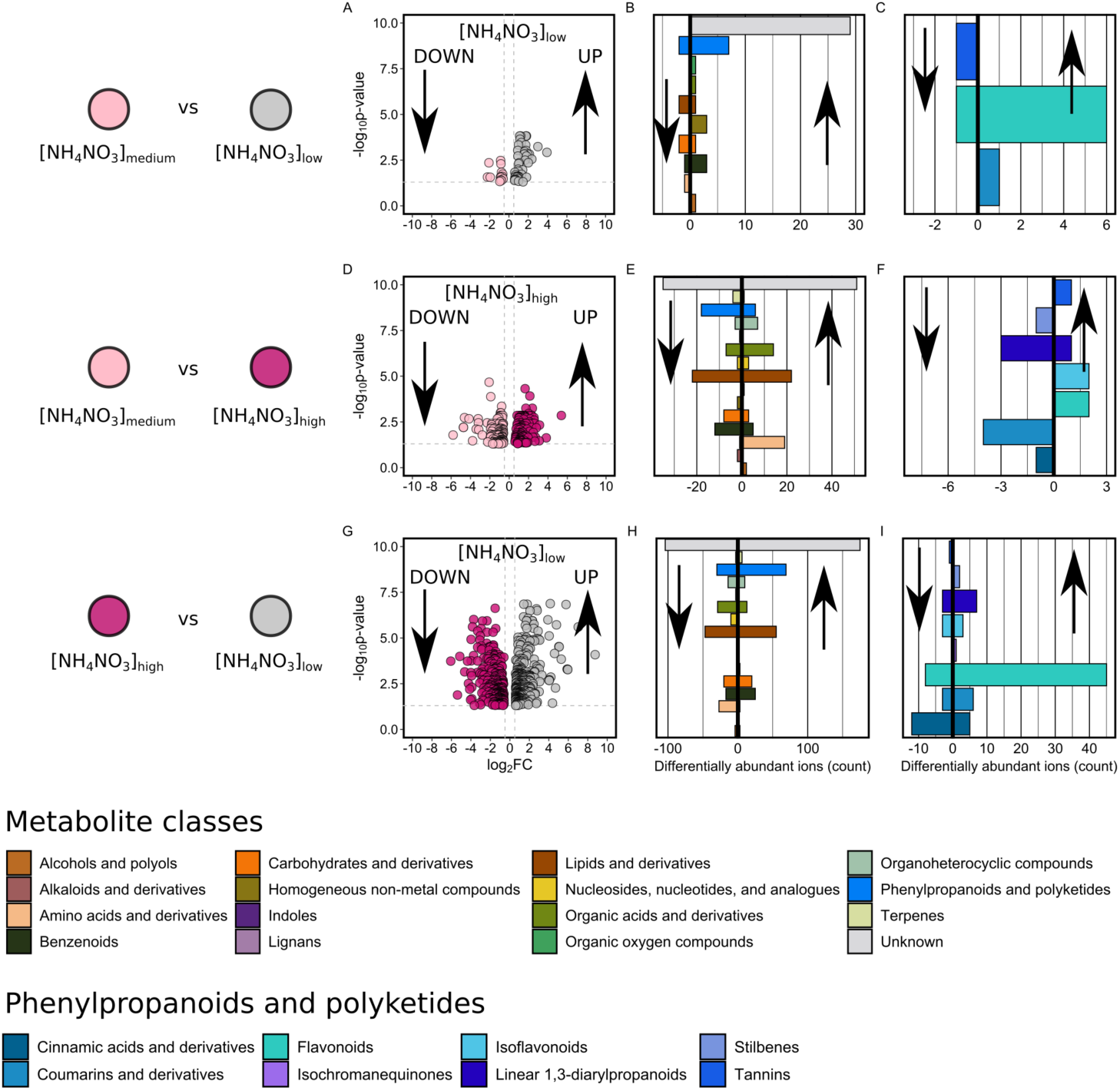
Quantitative differences in metabolite abundance between nitrogen levels in leaves from the early summer period (i.e. june-july) *(Vitis vinifera* cv. Sauvignon blanc). First line represents the comparison analysis between [NH4NO3]1ow and [NH4NO3]medium (A,B,C), second line represents the comparison between [NH4NO3]medium and [NH4NO3]high (D,E,F), and the third line represents the comparison analysis between [NH4NO3]high and [NH4NO3]1ow (G,H,1). (A,D,G) Volcano plots expressing statistical enrichment (“UP”) and impairment (“DOWN”) of features (Welch’s t-test) as a function of fold difference in each differential treatment comparison. Data shown represent negative (Electrospray ionization, ESI-) features from extractions. Cut-off values were set at P value < 0.05 and fold change > 1.5), using a Benjamini-Hochberg correction for false-discovery rate (FDR). (B,E,H) Class assignment of the putative metabolites selected through the volcano plots, either enriched (“UP” arrows) or impaired (“DOWN” arrows). Class assignment was performed using Classyfire and literature. Multiple features putatively annotating to the same metabolite were counted additively in the metabolite classes. Putative metabolites that are unlikely to accumulate as natural products (e.g. drugs) were classified as unknown along with feature markers that could not be assigned to any known compound. (C,F,I) Overview of the phenylpropanoids and polyketides class.

Comparing the [NH_4_NO_3_]_high_ level to the [NH_4_NO_3_]_medium_ level of the early summer period, we found 134 features significantly enriched in the high nitrogen level and 117 features significantly depleted (Fig. 8D). Among putative metabolites annotated, mainly amino acids, lignans, nucleosides, organic acids, organoheterocyclic compounds were enriched under [NH_4_NO_3_]_high_ level, while terpenes, phenylpropanoids, benzenoids and carbohydrates were globally depleted (Fig. 8E). Among the phenylpropanoids, we observed an over-representation in flavonoids, isoflavonoids, and tannins under [NH_4_NO_3_]_high_ level and an under-representation in diarylpropanoids, stilbenes, coumarines, and cinnamic acids (Fig. 8F).

Finally, comparing [NH_4_NO_3_]_low_ level to [NH_4_NO_3_]_high_ level in the early summer period, we found 390 features significantly enriched in the [NH_4_NO_3_]_high_ level and 311 features significantly depleted (Fig. 8D). Mainly benzenoids and phenylpropanoids were enriched and organic and amino acids depleted in [NH_4_NO_3_]_low_ level (Fig. 8H). The phenylpropanoids enrichment was driven by an over-representation in flavonoids (Fig. 8I).

The list of metabolic features significantly enriched or depleted in leaf samples for each of the above comparisons (Fig. 8) is presented in detail in Table S5.

According to the previously described results on the response of phenylpropanoids to nitrogen levels in the early summer period, we further explored their variability in the other period (summer; Supplementary Fig. S9; Table S6). We found that the difference in phenylpropanoids between [NH_4_NO_3_]_low_ level and [NH_4_NO_3_]_medium_ level or [NH_4_NO_3_]_high_ level was driven by an enrichment in flavonoids also in the summer period (Supplementary Fig. S9B,F). In contrast to the early summer period, flavonoids were enriched in the [NH_4_NO_3_]_medium_ level compared to [NH_4_NO_3_]_high_ level (Supplementary Fig. S9D). Regarding the trunk metabolome of control plants at the end of the season, PCA presented partial overlap between the three nitrogen levels (Fig. 7C). However, the discriminating volcano plot analysis showed 74 features enriched in [NH_4_NO_3_]_low_ level compared to [NH_4_NO_3_]_medium_ level and 77 depleted features (Supplementary Fig. S10A). Comparing the [NH_4_NO_3_]_high_ level to the [NH_4_NO_3_]_medium_ level, we found 124 features enriched and 117 features depleted (Supplementary Fig. S10C). In [NH_4_NO_3_]_low_ level compared to [NH_4_NO_3_]_high_ level, 196 features were enriched and 204 features were depleted (Supplementary Fig. S10E).

We found depleted levels of flavonoids and an enrichment in cinnamic acids in the [NH_4_NO_3_]_low_ level compared to [NH_4_NO_3_]_medium_ level. The differences between phenylpropanoids in the [NH_4_NO_3_]_medium_ and [NH_4_NO_3_]_high_ levels and between [NH_4_NO_3_]_low_ and [NH_4_NO_3_]_high_ levels were driven by the accumulation in flavonoid features in [NH_4_NO_3_]_medium_ level and [NH_4_NO_3_]_low_ level, respectively (Supplementary Fig. S10B,D).

The list of metabolic features significantly enriched or depleted in trunk samples for each of the above comparisons is presented in detail in Table S7.

Overall, metabolome analyses suggest a potential trade-off between primary and secondary metabolism depending on the level of stress (from deficit to excess) and the type of plant tissue (wood or leaf).

### Effect of nitrogen fertilization on grapevine susceptibility to wood pathogens

The necrosis sizes obtained following fungal inoculation were significantly higher than necrosis sizes obtained in negative controls (mock) for both pathogen species and the three nitrogen levels (P value < 0.001; Supplementary Fig. S11A,B,C). The necrosis caused by *N. parvum* inoculation did not differ significantly among the cuttings produced from the three nitrogen levels (Kruskal-Wallis test, p=0.92), and ranged from 0.5 to 3.9 cm, 0.9 to 4.5 cm, and 0.4 to 4.4 cm in [NH_4_NO_3_]low, [NH_4_NO_3_]medium and [NH_4_NO_3_]high, respectively (Supplementary Fig. S11A). The necrosis caused by *P. chlamydospora* also did not differ significantly among the cuttings produced from the three nitrogen levels (Kruskal-Wallis test, p=0.06), but tended to be longer in the low fertilization level. They ranged from 0.6 to 2.7 cm, 0.4 to 1.8 cm, and 0.9 to 1.2 cm in [NH_4_NO_3_]low, [NH_4_NO_3_]medium and [NH_4_NO_3_]high, respectively (Supplementary Fig. S11B).

## DISCUSSION

We investigated the effect of nitrogen deficiency or excess on esca disease incidence, fungal wood communities, and plant physiology and metabolism in *Vitis vinifera* cv. Sauvignon blanc over a three-year experiment. Consistent differences in N-tester index over the years and increasing vigor with increasing levels of nitrogen nutrition unequivocally confirmed the accuracy of the experimental design. Furthermore, the senescence process was advanced under nitrogen deficit and delayed under nitrogen excess, which is also well-documented in the literature (Agüera et al., 2010; Fan et al., 2023). Using this experimental design, we demonstrated that a deficit in nitrogen nutrition drastically reduced esca leaf symptom development, while an excess in nitrogen nutrition did not induce any significant difference compared to the medium nitrogen level. Through an integrative approach, we characterized plant physiological and biochemical changes with varying nitrogen fertilization, and demonstrated an absence of any impact on the wood fungal communities in control plants. This work provides new insights into the role of grapevine nutrition in the incidence of vascular disease and the putative underlying mechanisms.

### Impacts of nitrogen nutrition deficiency on esca incidence and plant physiology and metabolism

It is widely recognised that nitrogen availability has an influence on plant-pathogen interactions (Fagard *et al*., 2014). However, the nature of this impact remains complex and is driven by multiple factors such as the host and pathogen species studied and the type of nitrogen nutrition used. Here, we demonstrated that nitrogen deficiency led to a significant reduction in esca incidence.

Under nitrogen deficiency conditions, plant vigor and the average leaf area were reduced, as widely documented in the literature (Schreiner *et al*., 2013). We can therefore assume that nitrogen deficiency led to a reduction of the whole-plant (canopy) leaf area (as shown in sorghum (Zhao *et al*., 2005)). At the leaf level, we found no significant change in stomatal density. CO_2_ assimilation was reduced under nitrogen deficiency compared with plants under medium nitrogen nutrition. This result was expected since more than half of the nitrogen present in leaves is needed to synthesize the components of the photosynthetic apparatus (Evans, 1989). Nitrogen deficiency could induce a reduction in the activity and content of enzymes (such asRuBP and RuBISCO) and the concentration of chlorophyll (Dinu *et al*., 2022), which in turn negatively affects photosynthesis (Xie *et al*., 2020). Schreiner *et al*. (2013) also demonstrated that a reduction in nitrogen availability in Pinot Noir led to a decrease in CO_2_ assimilation. However, Squeri *et al*. (2021) who also worked on old potted vines (*Vitis vinifera* cv. ‘Barbera’), observed that photosynthesis was maintained under nitrogen deficit. Furthermore, we found that a decrease in nitrogen availability did not impact stomatal conductance when compared to the medium nitrogen level. Stomatal conductance has been shown not to be regulated under nitrogen deficiency (Keller *et al*., 1998). The reduced leaf area and vigor, without any modification in stomatal density or stomatal conductance, likely led to a reduction in transpiration at the whole-plant level.

It has been shown that reduced plant transpiration led to reduced esca leaf symptom incidence (Bortolami *et al*., 2021*b*), suggesting a key role of vine transpiration in esca pathogenesis. In addition, plant vigor has been hypothesized to be positively correlated with esca incidence in the vineyard but this hypothesis has rarely been tested. Only two recent studies suggested that cultivars characterized by low vigor also exhibited low esca incidence (Claverie *et al*., 2025; Gastou *et al*., 2024). Consequently, the reduced leaf symptom expression under nitrogen deficit in our study may be triggered by the reduction of transpiring area associated with the nitrogen deficiency treatment.

Nitrogen nutrition effects on plant disease expression may also be mediated by plant metabolome modifications (Fagard *et al*., 2014). The metabolome of control leaves varied significantly with nitrogen nutrition throughout the summer season. By comparing nitrogen deficiency with the medium nitrogen treatment in the early summer period, we showed a decrease in carbohydrates, lipids and amino acids, and an increase in phenylpropanoids and benzenoids. The decrease in primary metabolites in favor of secondary metabolites was accentuated during the summer period (data not shown). In Fritz *et al*. (2006), during nitrogen deficiency, the increase in phenylpropanoids was correlated with the induction of genes involved in their metabolism, such as phenylalanine ammonia lyase (PAL). Rossouw *et al*. (2017) also demonstrated that polyphenols and other products of the shikimate pathway were affected by nitrogen availability. Therefore, these metabolomic analyses point to a profound reorchestration of primary and secondary metabolisms in response to nitrogen levels, involving some compounds already reported in previous nutritional studies of plant-pathogen interactions. These results partly follow the carbon nutrient balance hypothesis (Bryant *et al*., 1983) and the growth differentiation balance hypothesis (Herms and Mattson, 1992), explaining that under limiting growth conditions, available resources would be more highly allocated to defence production. However, this increase in defence metabolites is usually associated with a maintenance of photosynthesis (Herms and Mattson, 1992). In our study, as previously discussed, we found a reduction in CO_2_ assimilation. However, in the case of drought stress, some studies demonstrated an activation of secondary metabolism even though photosynthesis was reduced (Ma *et al*., 2014). We can assume that the remaining CO_2_ assimilation was enough to support the production of secondary metabolites as the photosynthesis demand was likely reduced due to reduced growth. Fluxomics analyses could help address the flux imbalance between carbon and nitrogen pathways.

In our study, the increase in phenylpropanoids under nitrogen deficit was mainly due to an accumulation of flavonoids at the leaf level. An increase in flavonoids under nitrogen deficit has been described in numerous species (grapevine leaves (Squeri *et al*., 2021); broccoli leaves (Fortier *et al*., 2010); and tomato leaves (Stewart *et al*., 2001)). Flavonoids act as signals in plant-microorganism interactions and are characterised by their anti-fungal and antioxidant properties (Macheix *et al*., 2005; Treutter, 2006). Studies have shown that a nitrogen-triggered accumulation of soluble phenolic compounds can induce species-dependent resistance to pathogens (Leser and Treutter, 2005). In our study, we are interested in nitrogen stress-induced flavonoids, which can also be constitutively synthetised, but whose biosynthesis is enhanced or inhibited by nitrogen stress. The anti-pathogenic properties of flavonoids may be non-specific and result from their antioxidant properties. They could chemically process excess ROS generated by the plant during infection, but this is still a matter of debate (Hernández *et al*., 2009; Decros *et al*., 2019). Flavonoids are also involved in the hypersensitivity response, which is the primary defence mechanism of infected plants, and in programmed cell death (Treutter, 2006). An increase in these flavonoids during nitrogen deficiency could therefore prevent the development of symptoms by stimulating plant defences.

Similarly to leaves, nitrogen deficiency enhances plant defence secondary metabolites in wood. However, it was not triggered by flavonoids as shown in leaves. As the wood was sampled late in the season compared to leaf sampling, further research during the same period, earlier in the summer, should be conducted to get a better view of this secondary metabolism induction under nitrogen deficiency.

### Impacts of excess nitrogen nutrition on esca incidence and plant physiology and metabolism

In contrast to nitrogen deficiency, excess nitrogen had no significant impact on esca leaf symptom development compared to the medium nitrogen treatment. Many studies only included two different nitrogen levels (Keller *et al*., 2003; Squeri *et al*., 2021). However, it has been shown that physiological responses are not necessarily linear with nitrogen nutrition (Bondada and Syvertsen, 2003; Zheng *et al*., 2022). Excess nitrogen also had an impact on physiology compared with average nitrogen, but through a different process to that of nitrogen deficiency. Under nitrogen excess, the vigor of the plant increased, as well as the leaf surface area, without any change in stomatal density. The transpiring surface area was therefore increased, which might be expected to result in an increase in leaf symptoms compared to medium nitrogen level. However, this increase was compensated at leaf level by lower stomatal conductance, leading to a reduction in *CO_2_* uptake. This regulation enabled plants to increase their *WUEi.* We can therefore assume that the whole plant stomatal conductance was likely maintained, with consequently no significant differences with medium nitrogen level in terms of plant transpiration and leaf symptom incidence.

These physiological modifications are likely associated with altered plant metabolism. As such, we demonstrated a modification of the metabolome from high nitrogen level plants compared to medium nitrogen level. However at leaf scale, these modifications changed across the season. In early summer, phenylpropanoids were more abundant in control leaves and healthy wood from medium nitrogen levels than in those from high nitrogen levels, as opposed to mid-summer when the converse was true. This variable response was consistent with the absence of a significant difference in the incidence of esca symptoms between these two nitrogen levels.

### Nitrogen nutrition did not significantly modify the wood fungal communities in control plants

The main research studies assessing the impact of nitrogen fertilization on fungal communities have focused on soil communities (Ullah *et al*., 2019), in particular on microbial activity within the rhizosphere (Hester et al., 2018) and, more rarely, on leaf/shoot communities (Borruso *et al*., 2021). In our case, since the pathogens associated with esca are usually detected only in the trunk, we focused on the trunk fungal communities. We identified fungal taxa potentially associated with esca and other grapevine trunk diseases in the literature in all healthy (i.e. non necrotic) wood samples from symptomatic plants and control plants (except in two samples from the high nitrogen level). Our results showed that the wood fungal communities of control plants were not affected by nitrogen nutrition at the end of 2022 the season. We can hypothesize that either two seasons of differential nutrition was too short to modify the diversity and composition of the fungal communities, or that the trunk microbiota is relatively stable. Trunk microbiota has been shown to vary during the season (Bruez *et al*., 2014, 2020), but we can assume this would not therefore be due to variation in nitrogen availability. This is consistent with the fact that, in contrast to the large differences obtained in leaf metabolism between nitrogen levels during the growing season, differences in the metabolome of healthy wood were much less pronounced (Supplementary Fig. S10).

In addition, we found that fungal pathogen inoculations (*Neofusicoccum parvum* and *Phaeomoniella chlamydospora*) on rooted cuttings produced from the three nitrogen levels did not show any differences of necrotic lesion length. This is the first reported trial to suggest that nitrogen fertilization has no impact on Sauvignon blanc susceptibility to these two trunk pathogens or conversely on the fungal strains’ pathogenicity.

### Fungal communities in healthy wood partly differ between symptomatic and control plants

A significant difference in alpha diversity between symptomatic plants and control plants was only shown in the high nitrogen level. The abundance of wood pathogens typically associated with esca in the literature did not differ between asymptomatic and symptomatic plants as shown in other studies (Del Frari et al. 2019; Bruez et al. 2020), but we acknowledged the sample size of symptomatic plants was unfortunately low in the present study. The global fungal communities in healthy wood did not differ strongly between nitrogen levels and presence/absence of esca leaf symptoms. We might assume that nitrogen fertilization could impact pathogen activity and aggressiveness, although we were unable to demonstrate this through the inoculation of two pathogens in woody cuttings collected from the three different nitrogen levels. However, the wood metabarcoding approach was realised after the onset of symptom expression. Further research should investigate the impacts of nitrogen level and symptom formation on fungal communities across the whole vegetative season, especially prior to symptom formation and at the symptom onset, and on a large number of vines if possible. This would allow the identification of any possible shift in communities that could explain the development of symptoms and/or a possible imbalance within the communities depending on nitrogen level.

## CONCLUSION

To conclude, our three-year investigation into the effects of nitrogen fertilization on esca pathogenesis in Sauvignon blanc demonstrated a significant impact on leaf symptom expression. Our integrative approach revealed physiological and biochemical changes induced by nitrogen nutrition, likely underlying the observed esca variability. This study highlights that nitrogen nutrition could be a lever in managing the expression of foliar symptoms of vascular diseases as esca can be reduced under low fertilization. In addition, one of the key pieces of information that growers can gather from this experimental study is the fact that high fertilisation levels and high vigor are not increasing the risk of esca expression. Exploring the effects of different nitrogen nutrition methods in a range of cultivars and wine regions may provide new insights into the understanding of esca leaf symptom expression in the vineyard.

## ACKNOWLEDGEMENTS

We thank the experimental and administrative teams of UMR SAVE (INRAE, Bordeaux, France) for providing the materials, logistics, and assistance during the experiment. In particular we are very thankful to Marie Chambard and Mathéo Pinol Daubisse for assistance in gas exchange measurements and wood sampling. We thank Jérôme Jolivet, Sebastien Gambier (UMR SAVE) and Jean-Pierre Petit (UMR EGFV) for providing technical knowledge and support for plant transplantation and maintenance. We are grateful to Cédric Cassan (UMR BFP) for the extraction and LCMS analysis.

## AUTHOR CONTRIBUTIONS

N.D.A., C.E.L.D., G.A.G., designed the experiments; N.D.A., C.E.L.D., and N.F. conducted the esca symptom notations, nitrogen fertilization management, and wood sampling; N.D.A. and N.F. carried out the leaf samplings, measured vigor, leaf gas exchange and area; A.R., P.P. and the MetaboHUB-Bordeaux team carried out of untargeted metabolomic measurements (extraction and MS-DIAL analysis); G.C. realized the experiment of controlled inoculations of wood pathogens and analyzed the associated dataset; N.D.A. wrote the first draft of the article under the supervision of C.E.L.D. and G.A.G.; *all authors participated in scientific discussion regarding the experiments and methodologies used, edited, and agreed on the last version of the article*.

## DATA AVAILABILITY

All metabarcoding sequencing data are available under ENA accession number PRJEB85831. All the data supporting the findings of this study are available in the Supplementary Data of this article. Raw metabolomic and ecophysiological databases are available from the corresponding author upon reasonable request.

