## Supplementary figures for "Nitrogen nutrition impacts grapevine esca leaf symptom incidence, physiology and metabolism"

Supplementary Figures S1-S11 are available in this document. Supplementary Tables S1-S7 are included as a separate Excel file.

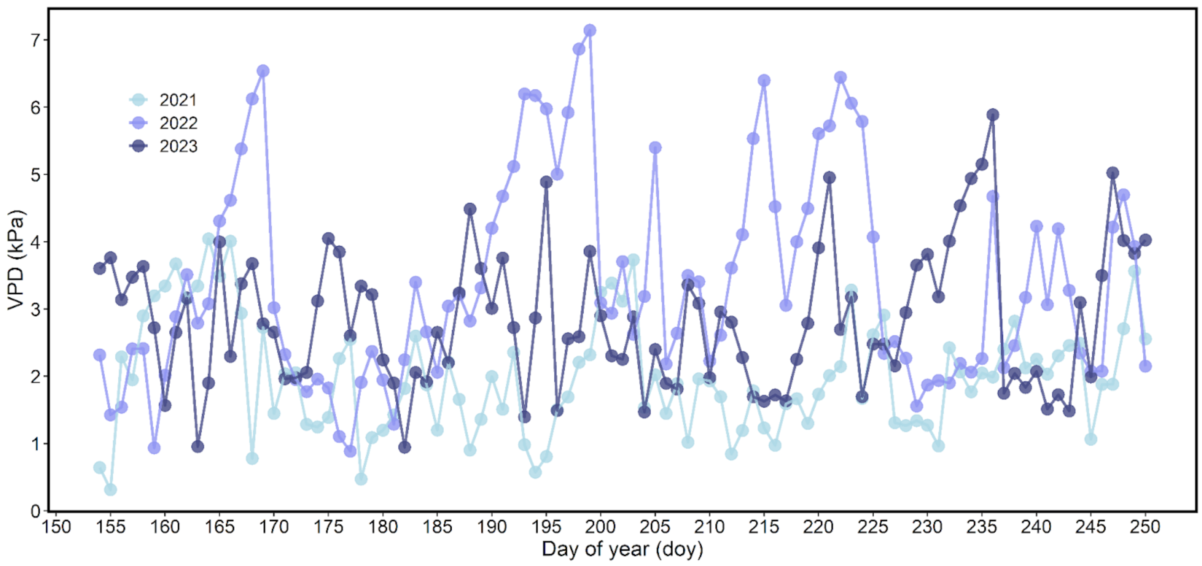


**Supplementary Figure S1. Evolution of the maximal Vapor Pressure Deficit (VPD, kPa) over time (day of year) during the three seasons of experiment.** The three years were 2021, 2022 and 2023 and are represented in light blue, light purple and dark purple respectively.

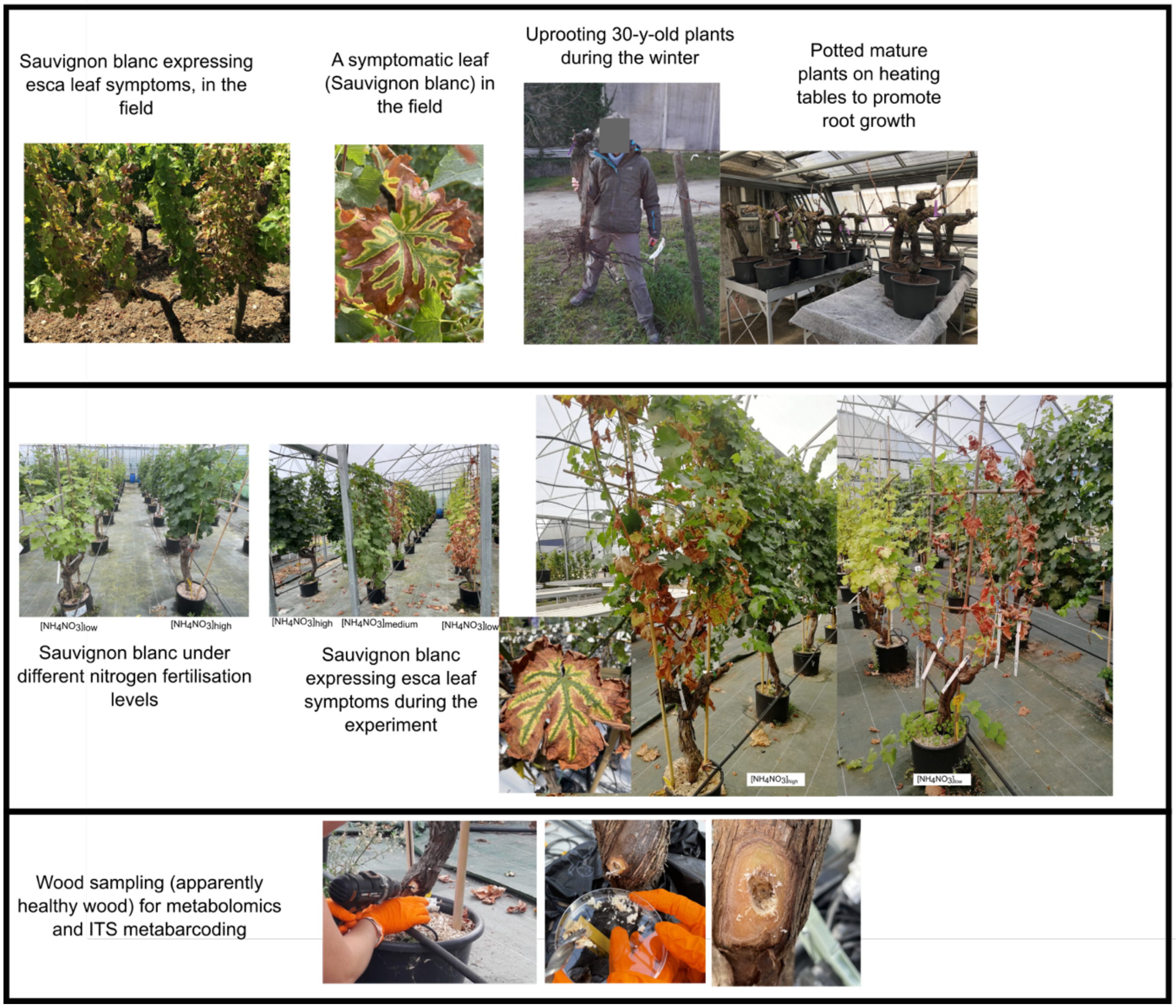


**Supplementary Figure S2. Images of plants (*Vitis vinifera* cv. Sauvignon blanc) and esca symptoms before, during and after the transplantation process, and sampling of apparently healthy wood using a drill.**

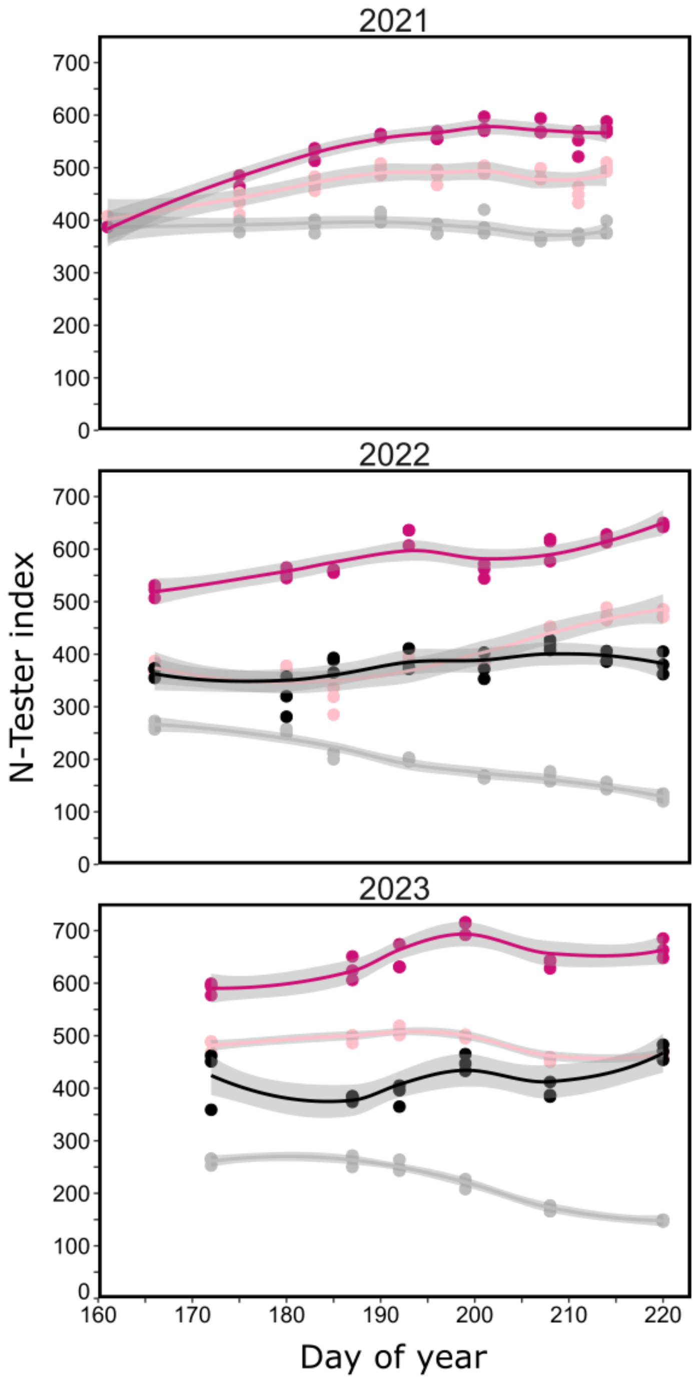


**Supplementary Figure S3. Evolution of the N-tester index over time (day of year) during the three seasons of experiment.** Low nitrogen level ($[NH_4NO_3]_{low}$, in grey), medium nitrogen level ($[NH_4NO_3]_{medium}$, in light pink) and high nitrogen level ($[NH_4NO_3]_{medium}$, in dark pink), and 'Sauvignon blanc' plants from the Vitadapt vineyard (in black) where measured in order to maintain the medium nitrogen level at the vineyard nitrogen level and the low and high nitrogen respectively above and under this target.

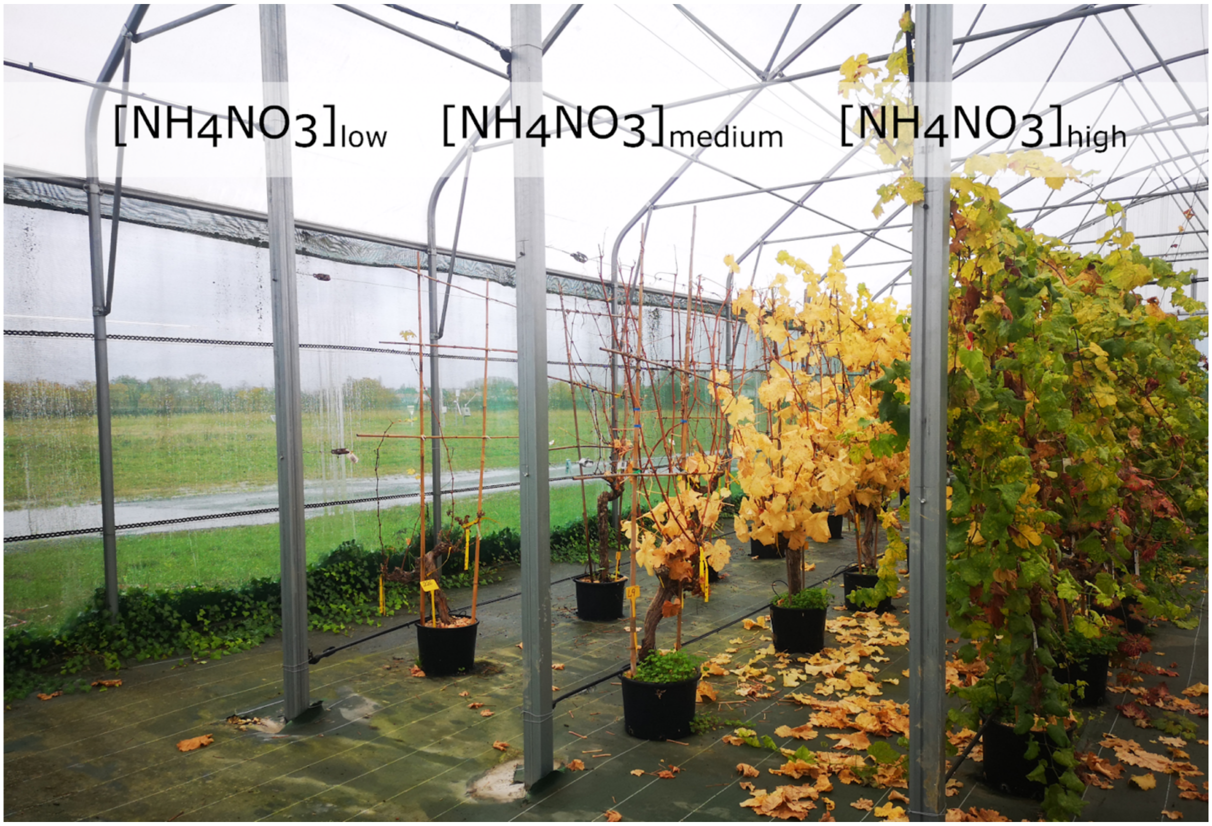


**Supplementary Figure S4. Picture from November 30, 2023 showing the differences in leaf senescence according to the three levels of nitrogen nutrition**: Low nitrogen level ($[NH_4NO_3]_{low}$,), medium nitrogen level ($[NH_4NO_3]_{medium}$) and high nitrogen level ($[NH_4NO_3]_{medium}$).

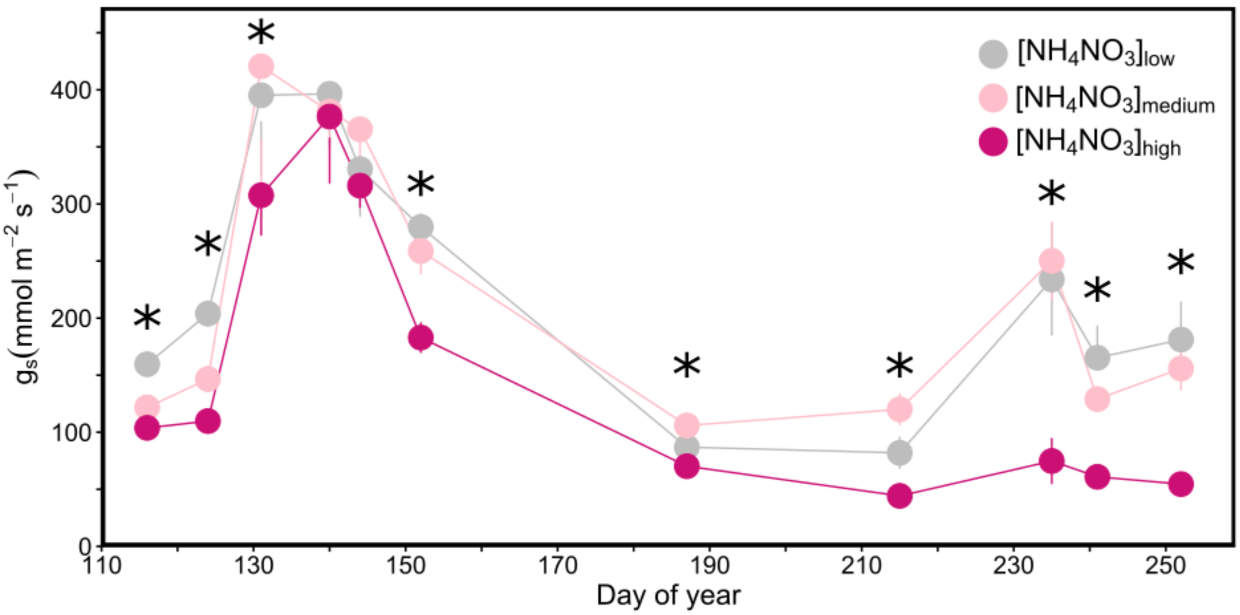


**Supplementary Figure S5. Leaf ambient stomatal conductance in control plants during the 2022 vegetative season.** Low nitrogen level ([NH4NO3]$_{low}$, 282 measurements in total) is represented in grey, medium nitrogen level ([NH4NO3]$_{medium}$, 250 measurements in total) in pink and high nitrogen level ([NH4NO3]$_{high}$, 289 measurements in total) in dark pink. Stars represent significant differences in ANOVAs ($P$ value < 0.05) between nitrogen levels for a specific date.

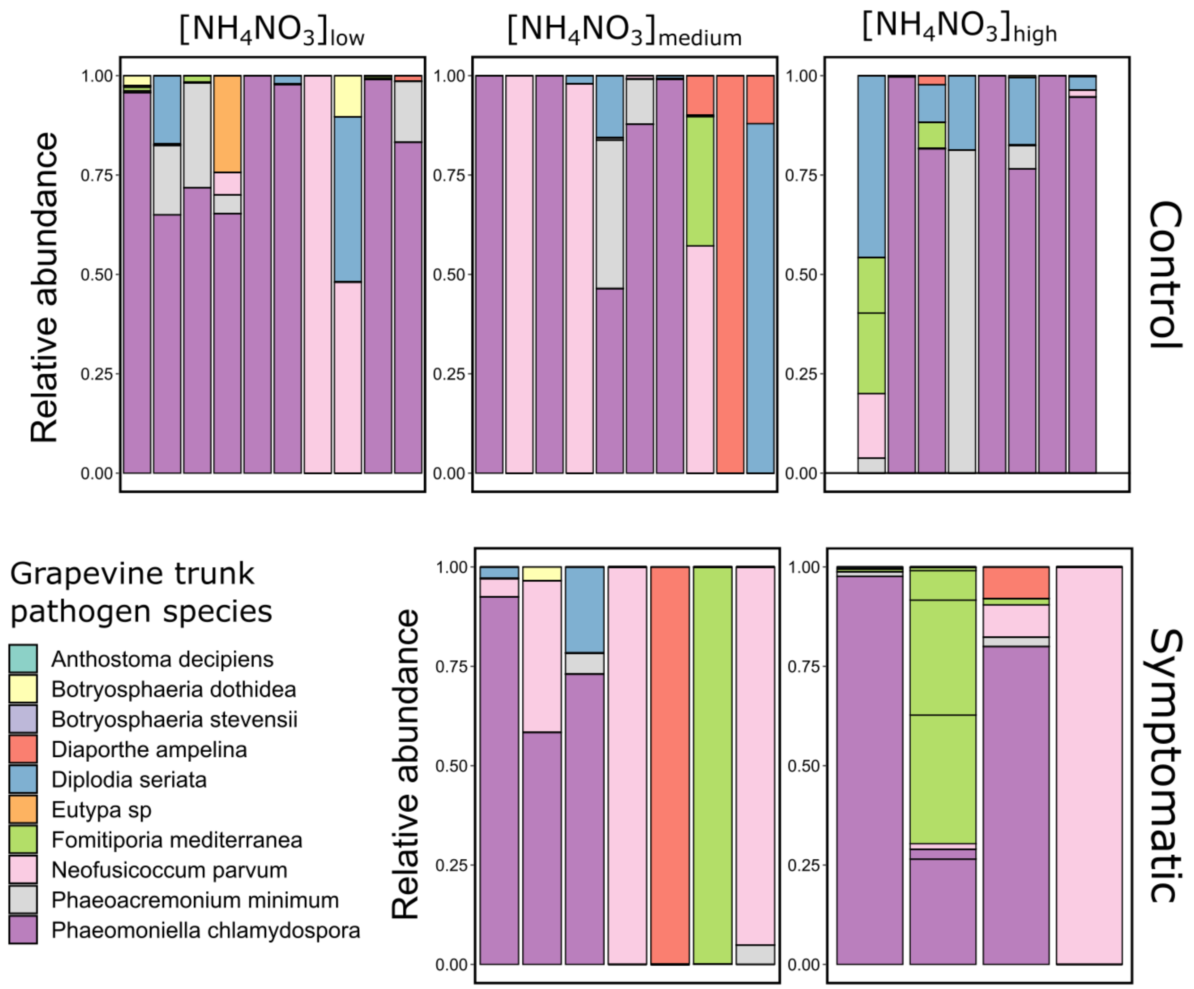


**Supplementary Figure S6. Relative abundance of the grapevine trunk pathogen species in healthy wood from control and symptomatic *Vitis vinifera* cv. 'Sauvignon blanc' from the three nitrogen levels.** Each bar represents one sample (n=41).

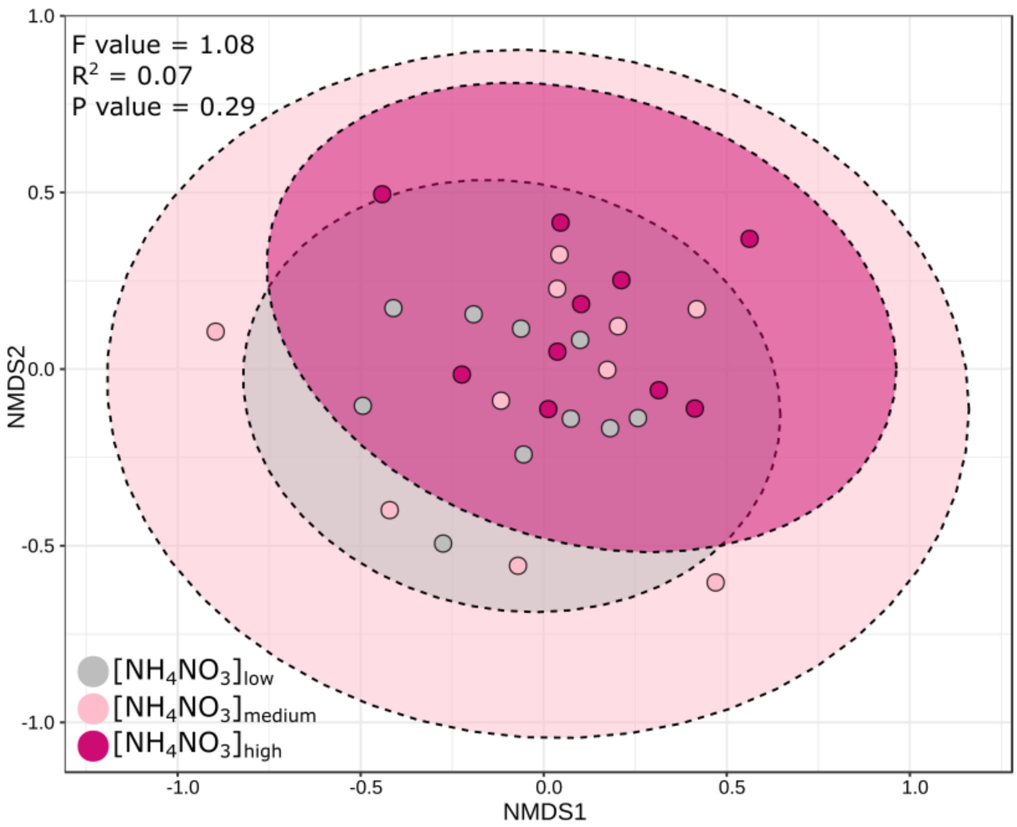


**Supplementary Figure S7. Ordination plot using a non-metric multidimensional scaling (NMDS) ordination matrix based on Bray-curtis distance comparing healthy wood samples from asymptomatic *Vitis vinifera* cv. Sauvignon blanc subjected to the three nitrogen levels**. Beta dispersion was calculated from transformed centred log-ratio (CLR) data to test the difference in dissimilarity between nitrogen levels in control plants using a PERMANOVA.

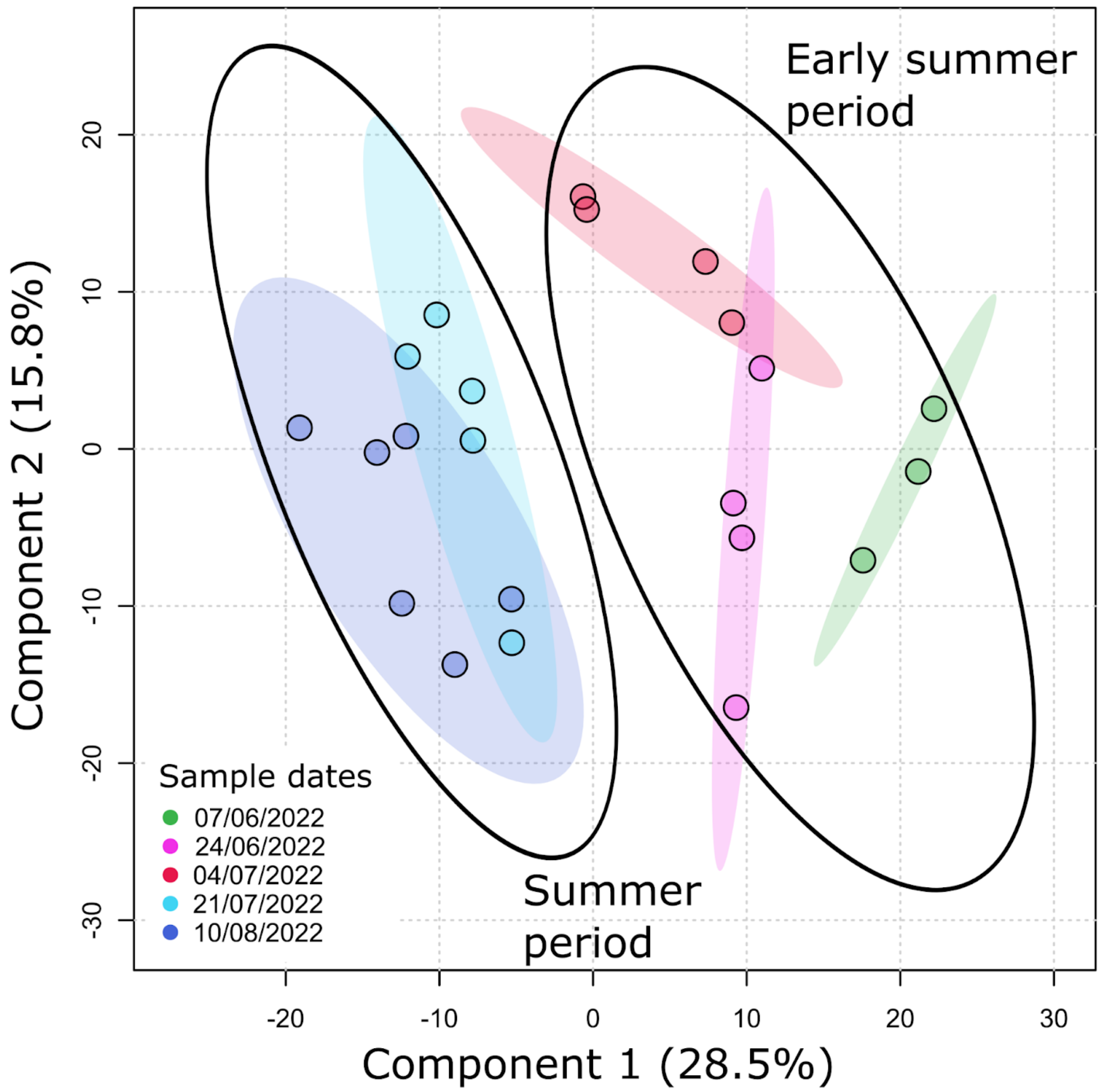


**Supplementary Figure S8. Principal component analysis (PCA) of metabolic profiles in control leaves from the medium nitrogen level during the season.** The early summer period includes dates between June 7, 2022 and July 4, 2022 and the summer period includes July 21, 2022 and August 10, 2022.

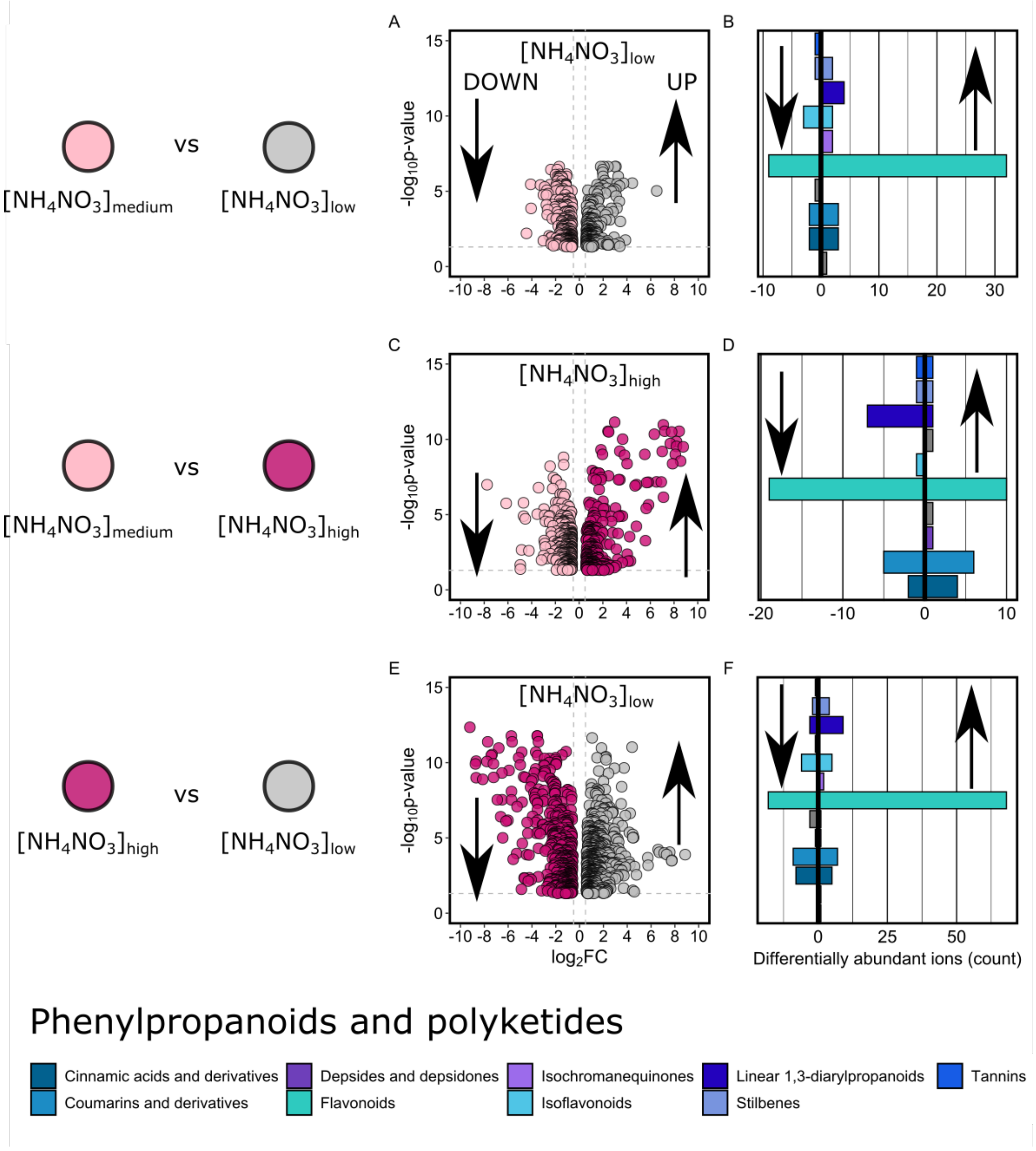


**Supplementary Figure S9. Quantitative differences in metabolite abundance (phenylpropanoids and polyketides class) between nitrogen levels in leaves from the summer period (July-August) (*Vitis vinifera* cv. 'Sauvignon blanc').** First line represents the comparison analysis between $[NH_4NO_3]_{low}$ and $[NH_4NO_3]_{medium}$ (A,B), second line represents the comparison between $[NH_4NO_3]_{medium}$ and $[NH_4NO_3]_{high}$ (C,D), and the third line represents the comparison analysis between $[NH_4NO_3]_{high}$ and $[NH_4NO_3]_{low}$ (E,F). (A,C,E) Volcano plots expressing statistical enrichment

(“UP”) and impairment (“DOWN”) of features (Welch’s t-test) as a function of fold difference in each differential treatment comparison. Data shown represent negative (Electrospray ionisation source, ESI–) features from extractions. Cut-off values were set at $P$ value < 0.05 and fold change > 1.5), using a Benjamini–Hochberg correction for false-discovery rate (FDR). (B,E,H) Class assignment of the putative metabolites selected through the volcano plots from the phenylpropanoids and polyketides class, either enriched (“UP” arrows) or impaired (“DOWN” arrows). Class assignment was performed using ClassyFire and literature. Multiple features putatively annotating to the same metabolite were counted additively in the metabolite classes. Putative metabolites that are unlikely to accumulate as natural products (e.g. drugs) were classified as unknown along with feature markers that could not be assigned to any known compound.

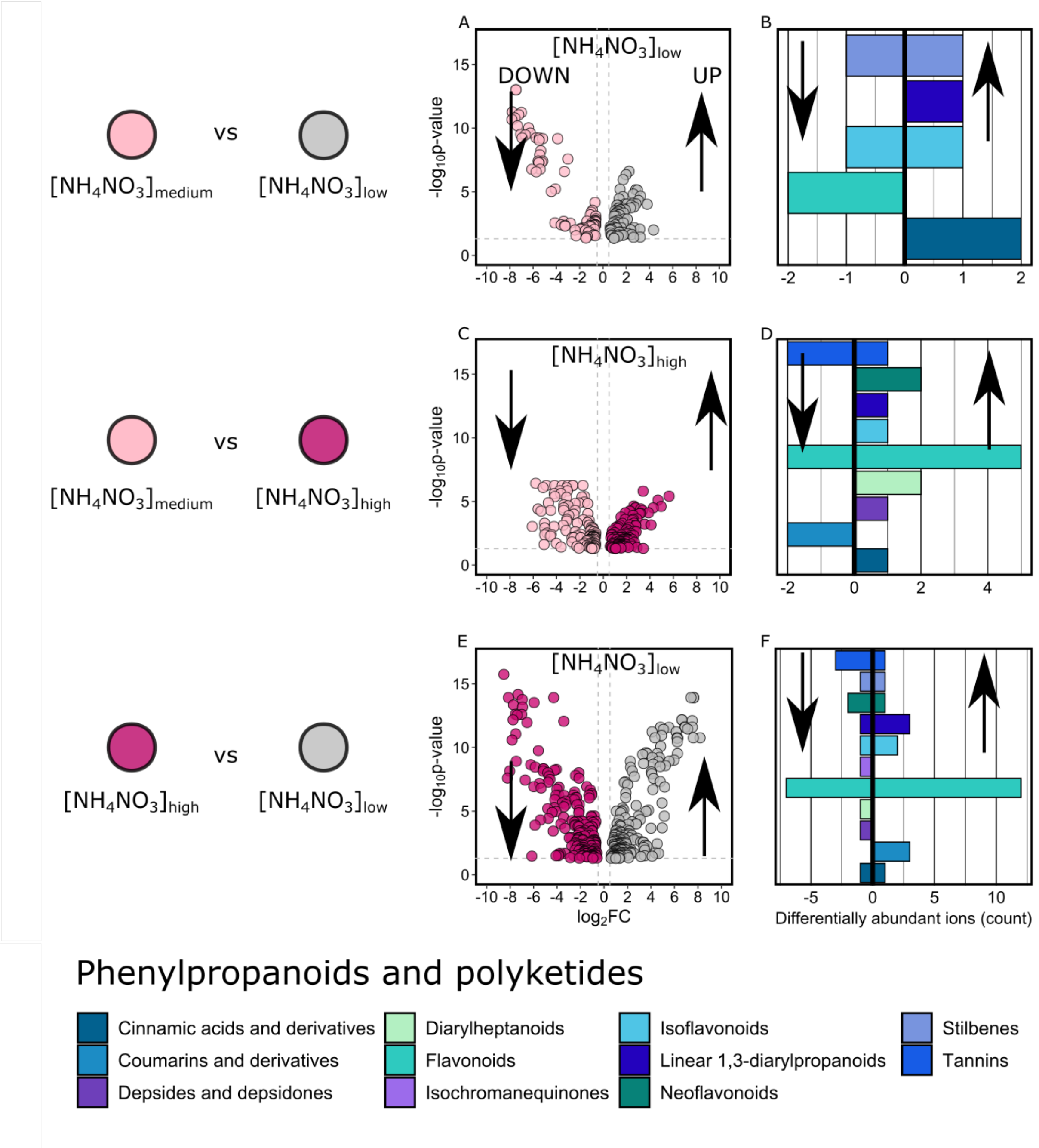


**Supplementary Figure S10. Quantitative differences in metabolite abundance (phenylpropanoids and polyketides class) between nitrogen levels in healthy wood from control *Vitis vinifera* cv. Sauvignon blanc.** First line represents the comparison analysis between $[NH_4NO_3]_{low}$ and $[NH_4NO_3]_{medium}$ (A,B), second line represents the comparison between $[NH_4NO_3]_{high}$ and $[NH_4NO_3]_{medium}$ (C,D) and the third line represents the comparison analysis between $[NH_4NO_3]_{low}$ and $[NH_4NO_3]_{high}$ (E,F). Volcano plots (A,C,E) expressing statistical enrichment ("UP") and impairment ("DOWN") of features (Welch's t-test) as a function of fold difference in differential treatments comparison: (A) enrichment ("UP") and impairment ("DOWN") features in $[NH_4NO_3]_{low}$ leaves compared to $[NH_4NO_3]_{medium}$ leaves, (C) enrichment ("UP") features and impairment ("DOWN") in $[NH_4NO_3]_{high}$ leaves compared to $[NH_4NO_3]_{medium}$ leaves, (E) enrichment ("UP") and impairment ("DOWN") features in $[NH_4NO_3]_{low}$ leaves compared to $[NH_4NO_3]_{high}$ leaves. Data shown represent negative (ESI–) features from extractions. Cut-off values were set at $P$ value < 0.05 and fold change > 1.5 using a Benjamini–Hochberg correction for false-discovery rate (FDR), (B,D,F) Class assignment of the putative metabolites selected through the volcano plots that are enriched ("UP") or impaired ("DOWN"). Class assignment was performed using ClassyFire and literature. Multiple features putatively annotating to the same metabolite were counted additively towards the metabolite classes. Putative metabolites that are unlikely to accumulate as natural products were added to the unknown metabolites that are feature markers that could not be assigned to any known compound.

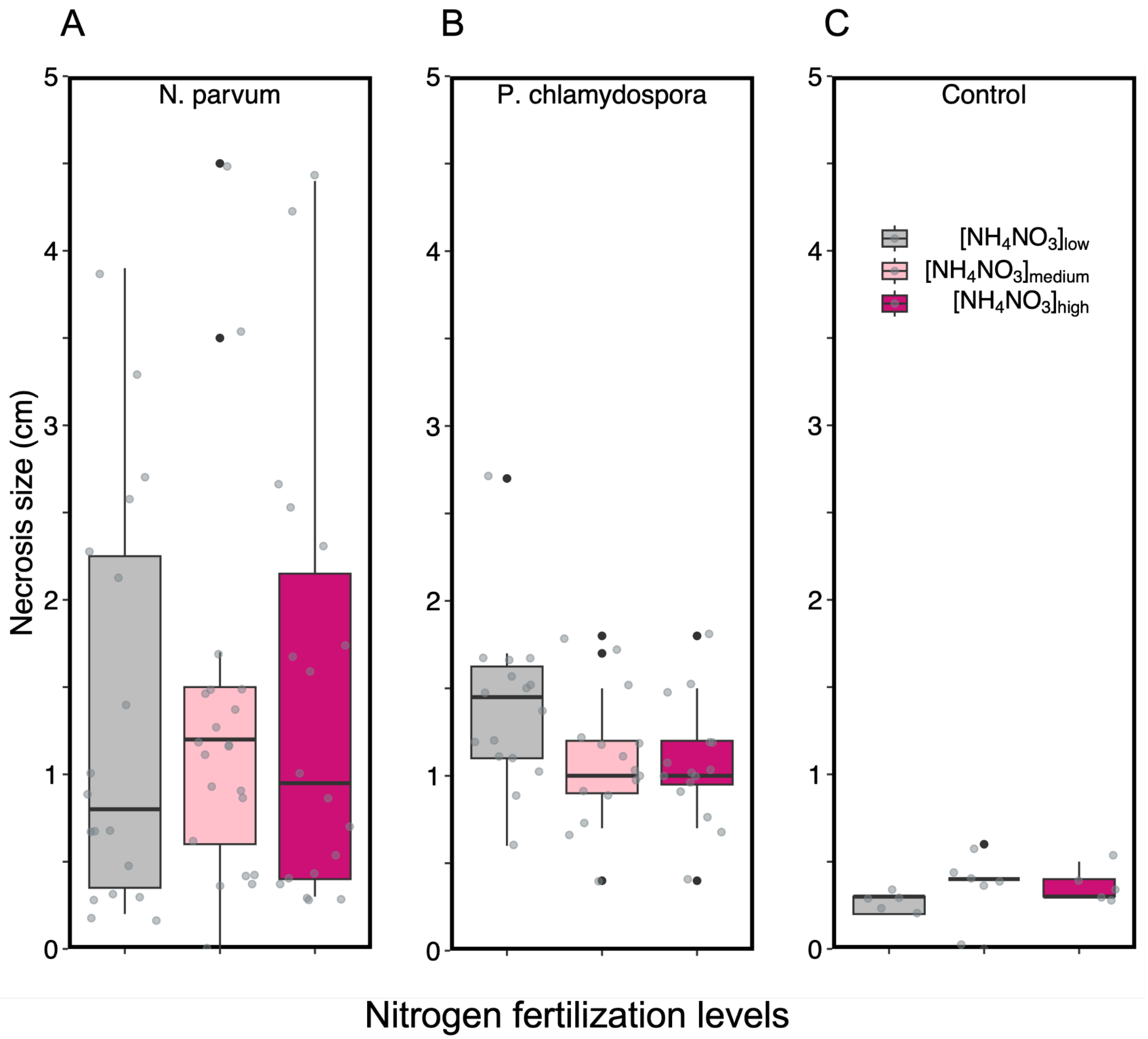


**Supplementary Figure S11. Necrosis sizes obtained following inoculation with *Neofusicoccum parvum* (A), *Phaeomoniella chlamydospora* (B), mock (control) inoculation (C) for each nitrogen level.** Fifteen internodes were inoculated for each nitrogen level and fungal species and 5 internodes were used as mock per nitrogen level. Boxplots display the median and interquartile range, with whiskers extending to the minimum and maximum values, excluding outliers, which are shown as individual black points. Individual data points are shown in grey.
